# Explainable Machine Learning Approach to Predict and Explain the Relationship between Task-based fMRI and Individual Differences in Cognition

**DOI:** 10.1101/2020.10.21.348367

**Authors:** Narun Pat, Yue Wang, Adam Bartonicek, Julián Candia, Argyris Stringaris

**Author notes:** Corresponding author: Narun Pat, PhD, also known as Narun Pornpattananangkul, Department of Psychology, University of Otago, William James Building, 275 Leith Walk, Dunedin 9016, New Zealand.

## Abstract

Despite decades of costly research, we still cannot accurately predict individual differences in cognition from task-based fMRI. Moreover, aiming for methods with higher prediction is not sufficient. To understand brain-cognition relationships, we need to explain how these methods draw brain information to make the prediction. Here we applied an explainable machine-learning (ML) framework to predict cognition from task-based fMRI during the n-back working-memory task, using data from the Adolescent Brain Cognitive Development (n=3,989). We compared nine predictive algorithms in their ability to predict 12 cognitive abilities. We found better out-of-sample prediction from ML algorithms over the mass-univariate and OLS multiple regression. Among ML algorithms, Elastic Net, a linear and additive algorithm, performed either similar to or better than non-linear and interactive algorithms. We explained how these algorithms drew information, using SHapley Additive explanation, eNetXplorer, Accumulated Local Effects and Friedman’s H-statistic. These explainers demonstrated benefits of ML over the OLS multiple regression. For example, ML provided some consistency in variable importance with a previous study (Sripada et al. 2020) and consistency with the mass-univariate approach in the directionality of brain-cognition relationships at different regions. Accordingly, our explainable-ML framework predicted cognition from task-based fMRI with boosted prediction and explainability over standard methodologies.

## Introduction

Task-based functional magnetic resonance imaging (fMRI) has been a prominent tool for neuroscientists since the early 90s (Kwong et al. 1992). One goal of task-based fMRI is to derive brain-based predictive measures of individual differences in cognitive abilities (Gabrieli et al. 2015; Dubois and Adolphs 2016). Yet, the goal of obtaining a robust, predictable brain-cognition relationship from task-based fMRI remains largely unattained (Elliott et al. 2020). Moreover, obtaining a method with a higher predictive ability may not be sufficient if we cannot explain how such a method draws information from different brain regions to make a prediction. Using an explainable machine-learning framework (Molnar 2019; Belle and Papantonis 2021), we aim at 1) choosing algorithms that extract information across brain regions to predict individual differences in cognition from task-based fMRI data with higher predictive ability and 2) explaining how these algorithms draw information to make the prediction. Ultimately, this explainable, predictive model can potentially be used in future studies that collect task-based fMRI to predict individual differences in cognitive abilities, especially if the model is built from well-powered data. This is similar to the use of polygenic scores in genetics studies (Torkamani et al. 2018).

Conventionally, to learn about which brain regions are associated with individual differences, neuroscientists use the mass-univariate approach (Friston 2007). Here, researchers first test many univariate associations between 1) fMRI BOLD at each brain region that varies as a function of task conditions (e.g., high vs. low working memory load) and 2) an individual-difference variable of interest (e.g., cognitive abilities). They then apply multiple comparison corrections, such as Benjamini-Hochberg’s false discovery rate (FDR) (Benjamini et al. 2001) or Bonferroni, to control for false conclusions based on multiple testing (Friston 2007). Accordingly, the mass univariate analyses allow for easy interpretation of the brain-cognition association at each brain region. However, this simplicity may come at a price. Recent findings have challenged the ability of the mass univariate approach in predicting individual differences (Kragel et al. 2021; Marek et al. 2022). For instance, in the context of resting-state fMRI and structural MRI, Marek and colleagues (2022), showed that mass univariate analyses had a much poorer ability in predicting individual differences in cognition, compared to multivariate analyses, or the techniques involving drawing information across different brain regions simultaneously in one model. Additionally, having separate tests for different brain regions in a mass-univariate fashion rests on the assumption that these regions are statistically independent of each other, which seems unrealistic given what is known about brain function. This focus on ‘marginal importance’ (i.e., the contribution from each brain region at a time, ignoring other regions), as opposed to ‘partial importance’(i.e., the contribution from multiple regions in the same model), further constrains our ability to achieve a holistic understanding of the relationship between the brain and individual differences (Chen et al. 2019; Debeer and Strobl 2020).

Multivariate analyses have been suggested as a potential solution to improve prediction and avoid the use of multiple comparison corrections (Chen et al. 2019; Kragel et al. 2020). The ordinary least squares (OLS) multiple regression is arguably the most widespread method for predicting a response variable from multiple explanatory variables simultaneously. For task-based fMRI, this means simultaneously having all brain regions as explanatory variables to predict a response variable of individual differences. In gist, the OLS multiple regression fits a plane to the data that minimizes the squared distance between itself and the data points (James et al. 2013; Kuhn and Johnson 2013). With uncorrelated explanatory variables, the OLS multiple regression has the benefit of being readily interpretable - each explanatory variable’s slope represents an additive effect on the response variable (Kuhn and Johnson 2013).

However, in a situation where strongly correlated explanatory variables are present, known as multicollinearity, the OLS multiple regression may misrepresent the nature of the relationship between brain activity and individual differences. For instance, with high multicollinearity, the OLS multiple regression can give very unstable estimates of coefficients and extremely high estimates of model uncertainty (reflected in large standard errors) (Graham 2003; Alin 2010; Monti 2011; P. Vatcheva and Lee 2016). Moreover, the direction of an explanatory variable’s coefficient may also depend on its relationship with other explanatory variables thus leading to sign flips (Courville and Thompson 2001; Beckstead 2012; Ray-Mukherjee et al. 2014). For example, when an explanatory variable has a strong, positive correlation with other explanatory variables and by itself has a weak, positive correlation with the response variable, the OLS multiple regression can unintentionally show a negative weight for this explanatory variable. Accordingly, it is crucial to examine this ‘suppression’ by comparing the directionality of the relationship estimated from the OLS multiple regression with those from the mass-univariate analyses (Conger 1974; Ray-Mukherjee et al. 2014).

To improve out-of-sample prediction and to, a certain extent, mitigate multicollinearity, many researchers have exchanged the classical OLS multiple regression for machine-learning (ML) algorithms (Dormann et al. 2013). Yet, it is still unclear how much improvement in prediction ML algorithms may lead to. Machine-learning algorithms include, but are not limited to, algorithms based on penalised regression (Kuhn and Johnson 2013), tree-based regression (Breiman et al. 2017) and Support Vector Machine (SVM) (Cortes and Vapnik 1995), such as Elastic Net (Zou and Hastie 2005), Random Forest (Breiman 2001), XGBoost (Chen and Guestrin 2016), and Support Vector Machine (SVM) with different kernels (Cortes and Vapnik 1995; Drucker et al. 1996). These algorithms usually require hyperparameters that can be estimated through a cross-validation (James et al. 2013). The algorithms differ in how the relationships between explanatory and response variables as well as among explanatory variables are modelled. Some algorithms, such as Random Forest, XGBoost and SVM with certain kernels (e.g., polynomial and Radial Basis Function, RBF) (Cortes and Vapnik 1995; Drucker et al. 1996) allow for non-linearity in the relationship between explanatory and response variables as well as interaction among different explanatory variables. Elastic Net, on the other hand, extends directly from the linear OLS multiple regression, but with an added ability to regularise the contribution of explanatory variables (James et al. 2013; Kuhn and Johnson 2013). Accordingly, Elastic Net is linear (i.e., assuming the relationship between explanatory and response variables to be linear) and additive (i.e., not automatically modelling interactions among explanatory variables). It is still unclear whether non-linear and interactive algorithms improve the predictive ability of task-based fMRI over linear and additive algorithms as well as over the OLS multiple regression and mass-univariate analyses. Therefore, it is important to compare the predictive performance across algorithms.

A major drawback of machine-learning algorithms used in the task-based fMRI (Kragel et al. 2020) is the difficulty to explain how each algorithm draws information from different brain regions in making predictions. Fortunately, recent developments in the explainable machine-learning framework have provided techniques that can improve explainability for many algorithms (Molnar 2019). Here we focused on four aspects of explainability. The first aspect is *variable importance*, or the relative contribution from each brain region when an algorithm makes a prediction. Linear and additive algorithms, such as Elastic Net and the OLS multiple regression, usually make a prediction based on a weighted sum of features. Accordingly, an explanatory variable with a higher coefficient magnitude has a higher weight in prediction, and thus the coefficient magnitude can readily be used as a measure of variable importance. For non-linear and interactive algorithms, such as Random Forest, XGBoost and SVM with polynomial and RBF kernels, the prediction is not made based on a weighted sum of features, and thus we need an additional ‘explainer’ to compute variable importance. SHapley Additive exPlanation (SHAP) (Lundberg and Lee 2017) is a newly developed algorithm-agnostic technique for variable importance. SHAP is designed to explain the contribution of each explanatory variable via Shapley values (Roth 1988). Based on Cooperative Game Theory, a Shapley value quantifies a fair distribution of compensation to each player based on his/her contribution in all possible coalitions where each coalition includes a different subset of players. When applying Shapley values to machine learning, researchers treat each explanatory variable as a player in a game, a predicted value as compensation and subsets of explanatory variables as coalitions. Shapley values reflect the weighted differences in a predicted value when each explanatory variable is included vs. not included in all possible subsets of explanatory variables. SHAP offers a computationally efficient approach for estimating Shapley values (Lundberg and Lee 2017). Using these measures of variable importance, one is able to demonstrate the similarity in contribution from each brain region based on different algorithms. Importantly, we would also be able to examine the consistency in variable importance across studies with similar fMRI tasks and individual difference variables.

The second aspect of explainability is *variable selection* (Heinze et al. 2018). There are existing statistical methods that could assist us further in selecting explanatory variables based on their variable importance and uncertainty around the variable importance, at least for some algorithms. For instance, for mass univariate analyses and the OLS multiple regression, a conventional p-value associated with each coefficient is often used for variable selection. For Elastic Net, a permutation-based approach called eNetXplorer (Candia and Tsang 2019) has recently been proposed. The central idea behind eNetXplorer is to fit two different sets of Elastic Net models, each set consisting of many realizations of n-fold cross-validated models. In the first set, Elastic Net models are fitted to predict the true response variable (target models), whereas in the second set, the models are fitted to predict a randomly permuted response variable (null models). For example, if the observations are participants in a study, then the target models will try to predict one participant’s response from the same participant’s set of explanatory variables (e.g., brain regions), whereas the null models will try to predict one participant’s response from another participant’s explanatory variable. Both the null and the target models are tuned and assessed via repeated cross-validation. Given that there is no relationship between the shuffled response and the predictors in the null models, any non-null predictive accuracy and coefficient estimates in the null models have to be spurious. Comparisons of the magnitude of the coefficient estimates in target models to null models, as well as of the frequency of feature selection in target models to null models, allow for near-exact inference for each individual explanatory variable (via the permutation tests) (Winkler et al. 2014; Helwig 2019). While eNetXplorer use cases were originally demonstrated for cellular and molecular “omics” data, this approach is widely applicable to many other scenarios aimed at uncovering predictors in a multi-variable setting. Indeed, there is a considerable overlap in the challenges faced in the analysis of omics and fMRI data (Antonelli et al. 2019), and as such, eNetXplorer may prove a valuable tool in the latter as well. With eNetXplorer, we can compare and contrast variable selection between the OLS multiple regression and Elastic Net with regards to Elastic Net hyperparameters, the uncertainty of the coefficients, coefficient magnitude after regularisation, and multicollinearity.

In addition to demonstrating relative contribution from different brain regions, the third aspect of explainability is to understand the extent to which predictive values from each algorithm change as a function of fMRI activity at these regions--in terms of the *pattern* (i.e., linearity vs. non-linearity) and *directionality* (i.e., positive vs. negative) of the relationship with the response variable. It is straightforward to examine the pattern and directionality of the univariate relationship for mass-univariate analyses. For instance, a “univariate effects” plot (Fox and Weisberg 2018) can show a linear fitted line between fMRI activity at each different region and their associated predicted values of the response variable. For multivariate algorithms, we need to consider the collinearity among explanatory variables that could distort the pattern and directionality of the influences from each explanatory variable (Molnar 2019). Accumulated Local Effects (ALE), a newly developed algorithm-agnostic explainer, is designed to help visualise how each explanatory variable in each algorithm impacts a predictive value on average (Apley and Zhu 2020). Importantly, ALE is specifically designed to handle data with moderate collinearity. With univariate effects and ALE plots, we could examine the similarity in pattern and directionality across algorithms. This can potentially reveal “suppression,” allowing us to check whether multivariate algorithms provide a similar directionality to the univariate algorithm.

The fourth aspect is to demonstrate the extent to which the variation of the prediction from these algorithms depends on the *interaction* of the explanatory variables. Some algorithms, including Random Forest, XGBoost and SVM with RBF and polynomial kernels, allow for interactions among explanatory variables. Friedman’s H-statistic (Friedman and Popescu 2008) is a metric of the interaction strength between each brain region and all other brain regions in predicting individual differences. In the case that interactive algorithms have higher predictive performance than additive algorithms, Friedman’s H-statistic can reveal interaction from certain brain regions that may account for the boost in prediction.

Our study used a large task-based fMRI dataset in children from the Adolescent Brain Cognitive Development (ABCD) study (Casey et al. 2018). We treated task-based fMRI activity during the working-memory ‘n-back’ task (Barch et al. 2013; Casey et al. 2018) as our explanatory variables. fMRI activity during the n-back task has been shown to correlate well with the performance of cognitive tasks in children and adults (Rosenberg et al. 2020; Sripada et al. 2020). For our response variables, we used individual differences in behavioural performance from 11 cognitive tasks and the g-factor, or the latent variable that captured the shared variance of behavioural performance across different cognitive tasks. Using the explainable machine-learning framework (Molnar 2019), we first identified algorithms that could achieve good accuracy in predicting individual differences from task-based fMRI data. Here we compared widely used nine algorithms: the mass univariate with the FDR and Bonferroni corrections, OLS multiple regression, Elastic Net, Random Forest, XGBoost and SVM with linear, polynomial and RBF kernels. We especially aimed to compare the predictive ability of linear and additive multivariate algorithms, including the OLS multiple regression and Elastic Net, against mass-univariate analyses and non-linear and interactive multivariate algorithms. We then applied various explainers (such as SHAP, eNetXplorer, ALE and Friedman’s H-statistic) to explain the extent to which these algorithms drew information from each brain region in making a prediction in four aspects: variable importance, variable selection, pattern and directionality and interaction. We focused on explaining the models predicting the g-factor, so that we could 1) examine if our framework can capture individual differences in cognitive abilities in general (i.e., not confined to specific cognitive tasks) and 2) compare variable importance in our study with that of a previous study (Sripada et al. 2020).

## Materials and Methods

### Data

We used the data from the ABCD Study Curated Annual Release 2.01 (Yang and Jernigan 2019). Our participants were 9-10-year-old children scanned with 3T MRI systems at 21 sites across the United States, recruited as reported previously (Garavan et al. 2018). Following exclusion (see below), there were 3,989 children (1,968 females). The ethical oversight of this study is detailed elsewhere (Charness 2018). The ABCD Study provided detailed data acquisition procedures and fMRI image processing that are also outlined in previous publications (Casey et al. 2018; Hagler et al. 2019).

### Explanatory Variables

The ABCD study applied Freesurfer to parcellate the brain based on Destrieux (Destrieux et al. 2010) and ASEG (Fischl et al. 2002) atlases. The parcellation resulted in data showing fMRI activity at 167 grey-matter (148 cortical surface and 19 subcortical volumetric) regions. We used fMRI activity at these 167 regions during the n-back task (Barch et al. 2013; Casey et al. 2018) as our explanatory variables. In this n-back task, children saw pictures featuring houses and faces with different emotions. Depending on the trial blocks, children needed to report whether the picture matched either: (a) a picture shown 2 trials earlier (2-back condition), or (b) a picture shown at the beginning of the block (0-back condition). We used fMRI measures derived from the [2-back vs 0-back] linear contrast (i.e., high vs. low working memory load), averaged across two runs.

### Response Variables

We tested the ability of fMRI during the n-back task to predict individual differences in cognitive abilities across all variables available in the dataset. First is the behavioural performance collected from the n-back task during the fMRI scan. Specifically, we used the accuracy of the 2-back condition as it is correlated well with the behavioural performance of other cognitive tasks collected outside of the scanner (Rosenberg et al. 2020). Nonetheless, predicting behavioural performance collected from the same fMRI task may have captured idiosyncratic variance that is specific to the task and session, not necessarily capturing individual differences in cognitive abilities per se. Thus, we also used behavioural performance from 10 cognitive tasks (Luciana et al. 2018; Thompson et al. 2019) collected outside of the fMRI session, as additional response variables.

Children completed the 10 ‘out-of-scanner’ cognitive tasks on an iPad during a 70-min inperson visit. A detailed description of these out-of-scanner cognitive tasks was provided elsewhere (Luciana et al. 2018). First, the Flanker task measured inhibitory control (Eriksen and Eriksen 1974). Second, the Card Sort task measured the cognitive flexibility (Zelazo et al. 2013). Third, the Pattern Comparison Processing task measured the processing speed (Carlozzi et al. 2013). Fourth, the Picture Vocabulary task measured language and vocabulary comprehension (Gershon et al. 2014). Fifth, the Oral Reading Recognition task measured language decoding and reading (Bleck et al. 2013). Sixth, the Picture Sequence Memory task measured the episodic memory (Bauer et al. 2013). Seventh, the Rey-Auditory Verbal Learning task measured auditory learning, recall and recognition (Daniel and Wahlstrom 2014). Eight, the List Sorting Working Memory task measured the working-memory (Bleck et al. 2013). Ninth, the Little Man task measured visuospatial processing via mental rotation (Acker and Acker 1982). Tenth, the Matrix Reasoning task measured visuospatial problem solving and inductive reasoning (Daniel and Wahlstrom 2014).

Lastly, in addition to these 11 response variables, we also derived a general factor of cognitive abilities, called the g-factor, from out-of-scanner cognitive tasks. Similar to previous work on fMRI and individual differences in cognitive abilities in adults and children (Dubois et al. 2018; Ang et al. 2020; Sripada et al. 2020), we applied a bifactor model of the g-factor using confirmatory factor analysis (CFA). Here we treated the g-factor as the general latent variable underlying performance across out-of-scanner cognitive tasks that is orthogonal to the three specific factors: language reasoning (capturing the Picture Vocabulary, Oral Reading Recognition, List Sorting Working Memory and Matrix Reasoning tasks), cognitive flexibility (capturing the Flanker, Card Sort and Pattern Comparison Processing tasks) and memory recall (capturing the Picture Sequence Memory and Rey Auditory Verbal Learning tasks). We applied robust maximum likelihood estimation (MLR) with robust (Huber-White) standard errors and scaled test statistics. To demonstrate model fit, we used scaled and robust comparative fit index (CFI), Tucker-Lewis Index (TLI), root mean squared error of approximation (RMSEA) with 90% CI of the g-factor. To implement the CFA, we used *lavaan* (Rosseel 2012) (version=.6-9) and *semPlot* (Epskamp 2015).

### Exclusion criteria

We followed the exclusion criteria recommended by the ABCD study (Jernigan 2019). We excluded data with structure MRI T1 images that were flagged as needing clinical referrals, having incidental findings (e.g., hydrocephalus and herniation) or not passing quality controls (including IQC_T1_OK_SER and FSQC_QC fields). Second, we excluded data with fMRI T2* images that did not pass quality controls or had excessive movement (dof > 200). Third, we excluded data flagged with unacceptable behavioural performance during the n-back task. Fourth, we removed all data from a Philips scanner due to a post-processing error in Release 2.01. Lastly, we identified outliers of the contrast estimates for each of the 167 regions using the 3 IQR rule and applied listwise deletion to remove observations with outliers in any region.

### Modelling pipeline

We first split the data into training (75%) and test (25%) sets. For machine-learning-based algorithms that needed hyperparameter tuning (see below), we implemented a grid search via 10-fold cross-validation within the training set. Here each algorithm created model candidates with a different combination of hyperparameters, and we considered the candidate model with the lowest root mean square error (RMSE) as the best model for each algorithm. To prevent data leakage, we fitted the CFA model of the g-factor to the observations in the training set and later applied this fitted model to the test set. Similarly, we separately applied the 3 IQR rule and data standardisation on the training and test sets.

To evaluate predictive performance, we used the test set and examined the similarity between predicted and observed (i.e., real) values of each response variable. To reveal different aspects of the model’s predictive ability, we used multiple out-of-sample prediction metrics (Poldrack et al. 2020).

The first metric is Pearson’s correlation, which was defined as

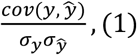

where *cov* is the covariance, *σ* is the standard deviation, *y* is the observed value and 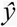 is the predicted value. Pearson’s correlation ranges from −1 to 1. The high positive Pearson’s correlation reflects high predictive accuracy, regardless of scale. Negative Pearson’s correlation reflects poor predictive information in the model.

Second, we used traditional *r* square defined using the sum-of-squared formulation:

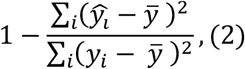

where 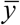 is the mean of the observed value. Traditional *r* square is often interpreted as variance explained, with the value closer to 1 reflecting high predictive accuracy. Like Pearson’s correlation, traditional *r* square can be negative in case of no predictive information in the model.

Third, we defined the mean absolute error (MAE) as

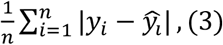

MAE measures how far the predicted value is from the observed value in absolute terms. Unlike Pearson’s correlation and traditional r square, MAE does not scale the data, rendering it more sensitive to scaling. Lower MAE reflects high predictive accuracy.

Forth, we defined the root mean square error (RMSE) as

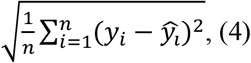

Similar to MAE, RMSE measures how far the predicted value is from the observed value but uses the square root of the average of squared differences, as opposed to the absolute differences. Lower RMSE reflects high predictive accuracy.

To demonstrate the distribution of the predictive performance across algorithms, we bootstrapped these out-of-sample prediction metrics on the test set 5000 times, resulting in bootstrap distributions of each prediction metric for each algorithm. To compare predictive performance, we also created bootstrap distributions of the differences in the predictive performance between each pair of algorithms. If the 95% confidence interval of the bootstrap distribution of the differences did not include zero, we concluded that the two algorithms were significantly different from each other. We used ‘tidymodels’ (www.tidymodels.org) for this pipeline.

### Modelling algorithms

We tested the predictive performance of nine algorithms. The first algorithm was the mass univariate analyses with the false discovery rate (FDR) correction. Here we used the simple linear regression with each of 167 regions as the only explanatory variable in each model, resulting in 167 different models in the form of:

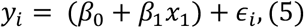

where *x* is the explanatory variable, *β* is the coefficient, and *ϵ* is the error term. *β* is estimated based on the minimization of the sum of squared errors following:

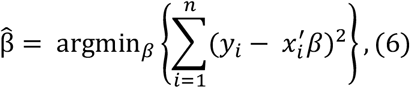

Where *n* is the number of observations (i.e., participants) in the training set. The FDR (Benjamini and Hochberg 1995) corrects for multiple comparisons following:

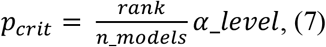

where *p_crit_* indicates a criterion in which p-value needs to be below, *rank* is the rank of p-values of one model compared with other models whereas the smallest p-value leads to *rank* = 1, *n_models* is the number of models, which equals 167 brain regions in our case, and α_level is the overall type-I error rate, set at .05. We then used the test set to examine the prediction of the models that passed the FDR correction in the training set at *p_crit_*.

The second algorithm was the mass univariate analyses with the Bonferroni correction. The Bonferroni correction is usually considered more conservative than the FDR correction (Moran 2003). We used the same simple regression with the first algorithm, except that here we applied the Bonferroni correction instead of the FDR correction. The Bonferroni correction defines *p_crit_* as

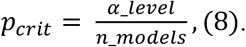

Note, only the first two algorithms were univariate algorithms, using separate models for each explanatory variable. Other algorithms were multivariate algorithms that included all brain regions as explanatory variables in one model. Accordingly, when creating bootstrap distributions of each prediction metric for univariate algorithms in the test set, we included multiple predicted values per testing participant based on the number of brain regions that survived each multiple comparison correction in the training set. By contrast, bootstrap distributions of each prediction metric for multivariate algorithms in the test set were based on one predictive value per testing participant for each algorithm.

The third algorithm was the ordinary least-square (OLS) multiple regression. Here, similar to the mass univariate analyses, the OLS multiple regression uses the same minimization of the sum of squared errors as the first two algorithms. However, the OLS multiple regression used all brain regions as explanatory variables in one linear regression model in the form of

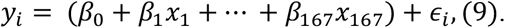

Note, all other algorithms below beyond the first three were machine-learning based that required hyper-parameter tuning via cross-validation.

The fourth algorithm was Elastic Net (Zou and Hastie 2005). Here, apart from minimizing the sum of squared errors done in the previous three algorithms (see equation #6), Elastic Net simultaneously minimises the weighted sum of the explanatory variables’ coefficients (Zou and Hastie 2005; James et al. 2013; Kuhn and Johnson 2013). As a result, Elastic Net shrinks the contribution of some explanatory variables closer towards zero (or set it to zero exactly). The degree of penalty to the sum of the explanatory variable’s coefficients is determined by a ‘penalty’ hyperparameter denoted by λ. The greater the penalty, the stronger shirking the explanatory variable’s coefficient is, and the more regularised the model becomes. In addition to the ‘penalty’ hyperparameter, Elastic Net also includes a ‘mixture’ hyperparameter denoted by a, which determines the degree to which the sum of either the squared (known as ‘Ridge’) or absolute (known as ‘LASSO’) coefficients is penalized. The estimates from Elastic Net are defined by

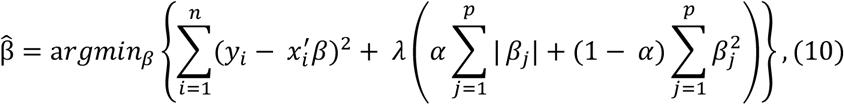

where *p* is the number of parameters. In our hyperparameter-turning grid, we used 200 levels of penalty from 10^-10 to 10, equally spaced on the logarithmic-10 scale and 11 levels of the mixture from 0 to 1 on the linear scale.

The fifth algorithm is Random Forest (Breiman 2001). Random Forest creates several regression trees by bootstrapping observations and including a random subset of explainable variables at each split of tree building. To make a prediction, Random Forest applies ‘bagging,’ or aggregating predicted values across bootstrapped trees. Unlike the above algorithms, Random Forest allows for 1) non-linearity (i.e., the relationship between explanatory and response variables does not constrain to be linear) and 2) interaction (i.e., different explanatory variables can interact with each other). Here we used 500 trees and tuned two hyperparameters: ‘mtry’ and ‘min_n’. ‘mtry’ is the number of explainable variables that are randomly sampled at each split. Here we used the integers between 1 to 167 (i.e., the maximum number of brain regions) in our grid. ‘min_n’ is the minimum number of observations to allow for a split of a node. In other words, min_n puts a limit on how trees grow. We used the integers between 2 to 2000 for min_n in our grid.

The sixth algorithm is XGBoost (Chen and Guestrin 2016). XGBoost is another regression-tree based algorithm. Like Random Forest, XGBoost also allows for non-linearity and interaction. Unlike ‘bagging’ used in Random Forest that builds multiple independent trees, ‘boosting’ used in XGBoost creates sequential trees where a current tree adapts from previous trees. Accordingly, XGBoost has ‘learning rate’ as a hyperparameter, denoted by *η*, to control for the speed of the adaptation. Here we sampled ‘learning rate’ from exponentiation with a base of 10 and 3,000 exponents ranging linearly from −10 to −1. XGBoost’s trees are also different from regular regression trees in many aspects. They, for instance, require a ‘loss reduction’ hyperparameter, denoted by γ, to control for the conservativeness in tree pruning. We sampled ‘loss reduction’ from exponentiation with a base of 10 and 3,000 exponents ranging linearly from −10 to 1.5. We also tuned ‘tree_dept’, or the maximum splits possible using integers from 1 to 15. Additionally, we tuned ‘sample_size’ or the proportion of observations exposed to the fitting routine, using 3,000 numbers ranging from .1 to .99 on the linear scale. Similar to Random Forest, here we used 500 trees and tuned ‘mtry’ from 1 to 167 and ‘min_n’ from 2 to 1000. To cope with many hyperparameters, we used a Latin hypercube grid with a size of 3,000.

The seventh to ninth algorithms are Support Vector Machine (SVM) for regression, also known as Support Vector Regression, with three different kernels: linear, polynomial and Radial Basis Function (RBF) (Cortes and Vapnik 1995; Drucker et al. 1996). SVM includes a ‘margin of tolerance’ hyperparameter, denoted by *ε*. The ‘margin of tolerance’ signifies an area around a hyperplane where no penalty is given to errors following:

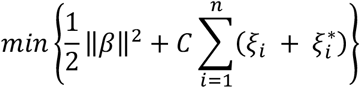

with constraints:

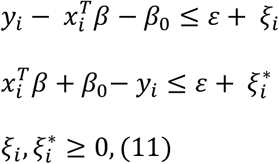

where *ξ* are non-zero slack variables that are allowed to be above (*ξ_i_*) and below 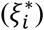 the margin of tolerance, and *C* is a ‘cost’ hyperparameter. A higher ‘cost’ indicates higher tolerance to data points outside of the ‘margin of tolerance’. Here we sampled the ‘margin of tolerance’ using 30 numbers from 0 to .2 on the linear scale. As for the ‘cost’, we sampled it from exponentiation with a base of 10 and 15 exponents ranging linearly from −3 to 1.5. To allow for non-linearity and interaction, researchers can apply kernel tricks to transform the data into a higher-dimensional space. Accordingly, in addition to using a linear kernel, we also applied polynomial and RBF kernels. For the polynomial kernel, we needed to tune two additional hyperparameters: ‘degree’ and ‘scale factor’. We sampled ‘degree’ from the numbers 1, 2 and 3 and ‘scale factor’ from exponentiation with a base of 10 and 10 exponents, ranging linearly from −10 to −1. As for the RBF kernel, we sampled another hyperparameter ‘RBF sigma’ from exponentiation with a base of 10 and 10 exponents, ranging linearly from −10 to 0.

### Explaining the algorithms

We applied different methods to help explain how each algorithm drew information from 167 brain regions when making predictions (Molnar 2019). We focused our analyses on the g-factor, so that we can compare our modelling explanation with prior findings with a similar exploratory and response variable in adults (Sripada et al. 2020).

#### Variable importance: Coefficients and SHapley Additive explanation (SHAP)

Variable importance refers to the relative contribution of each explanatory variable (i.e., brain region) in making a prediction. To better understand the similarity in contribution from each brain region based on different algorithms, we calculated Spearman’s correlation between variable importance of different algorithms. For the mass univariate analyses, the OLS multiple regression and Elastic Net, we used the coefficient magnitude as a measure for variable importance. Note for the mass univariate analyses, we did not apply any multiple-comparison correction when examining Spearman’s correlations in variable importance with other algorithms, so that the number of explanatory variables stayed the same across algorithms. For Random Forest, XGBoost and SVM, we applied SHapley Additive exPlanation (SHAP) (Lundberg and Lee 2017), to compute variable importance. We implemented SHAP using the *fastshap* package (https://bgreenwell.github.io/fastshap/).

In addition to examining the similarity in variable importance across algorithms, we also tested how variable importance found in the current study was in line with prior findings (Sripada et al. 2020). Sripada and colleagues (2020) have recently used young adult data from the Human Connectome Project (HCP) dataset and examined the multivariate relationship between task-fMRI from the similar n-back task and the g-factor. They used an algorithm based on the Principal Component Regression (PCR) and regressed the g-factor on the first 75 principal components (PCs). To explain the regions that were related to the g-factor, they projected all PCs back to the brain space, weighted by the magnitude of the regression. Fortunately, Sripada and colleagues (2020) uploaded this weighted, PCR brain map on https://balsa.wustl.edu/study/v0D7.

We downloaded and parcellated Sripada and colleagues’ (2020) brain map using Destrieux’s for cortical surface and ASEG for subcortical regions. We then examined Spearman’s correlations between our variable importance from different algorithms and Sripada and colleagues’ (2020) weighted, PCR brain map. Given the differences in the age of ABCD vs. HCP participants and the superiority of cortical surface in brain registration across ages (Ghosh et al. 2010), we separately computed Spearman’s correlations on the cortical surface alone and on the whole brain (including both cortical surface and subcortical volume).

#### Variable selection for variable importance: Conventional p-value and eNetXplorer

Here, we only focused on variable selection methods designed for algorithms based upon the general linear model, including mass univariate analyses, the OLS multiple regression and Elastic Net. We examined the variable selection of these algorithms using the training set. With the mass univariate analyses, we determined brain regions that were significantly associated with the g-factor using FDR and/or Bonferroni corrections (α-level = .05). With the OLS multiple regression, we defined significant regions as those with a p-value < .05 coefficient.

With Elastic Net, we selected the best mixture hyperparameters from the previously run grid search and applied eNetXplorer (Candia and Tsang 2019) to fit two sets of many Elastic Net models. In one set, the models were fitted to predict the true response variable (the g-factor; target models), while in the other set the models were fitted to predict the same response variable randomly permuted (null permuted models). We split the data into 10 folds 100 times (100 runs; eNetXplorer default), and then in each run, the target models were repeatedly trained on 9 folds and tested on the leftover fold. Additionally, for each cross-validation run of the target models, there were 25 permutations of the null permuted models (eNetXplorer default). We defined the explanatory variables’ (brain regions) coefficient for each run, *β^r^*, as the average of non-zero model coefficients across all folds in a given run. Across runs, we used an average of a model coefficient weighted by the frequency of obtaining a non-zero model coefficient per run. See the detailed implementation of eNetXplorer (Candia and Tsang 2019). Formally, we defined an empirical p-value as:

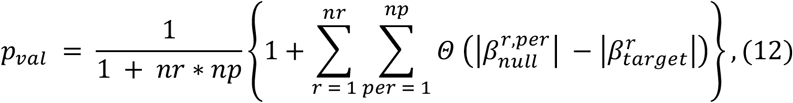

where *p_val_* is an empirical p-value, *r* is a run index, *nr* is the number of runs, *per* is a permutation index, *np* is the number of permutations, Θ is the right-continuous Heaviside step function and |β| is the magnitude of an explanatory variable’s coefficient. That is, eNetXplorer uses the proportion of runs in which the null models performed better than the target models to establish statistical significance for each explanatory variable’s coefficient. We implemented eNetXplorer using the *eNetXplorer* package version 1.1.2 (https://github.com/cran/eNetXplorer).

To demonstrate the differences in variable selection of the OLS multiple regression and eNetXplorer, we created plots to visualise coefficient magnitude and uncertainty estimates as a function of hyperparameters of Elastic Net and multicollinearity. For uncertainty estimates, we used standard error (SE) of the coefficients for the OLS multiple regression and standard deviation (SD) of the permuted null coefficients from eNetXplorer for Elastic Net. As for hyperparameters of Elastic Net, we created three plots with different solutions: full Elastic Net (i.e., tuning both the ‘mixture’ (α) and ‘penalty’ (λ) hyperparameters), Ridge (i.e., fixing the mixture at 0 and tuning the penalty) and LASSO (i.e., fixing the mixture at 1 and turning the penalty). Lastly, we quantified multicollinearity of the brain regions using Variance Inflation Factors (VIF) calculated from the model fit of the unregularized model (i.e., the OLS multiple regression).

#### Prediction pattern and directionality: Univariate Effects and Accumulated Local Effects

Here we demonstrated the pattern (i.e., linearity vs. non-linearity) and directionality (i.e., positive vs. negative) of the relationship between task-based fMRI activity at different brain regions and the g-factor based on different algorithms. For mass-univariate analyses, we plotted a ‘univariate effect’, a fitted line between fMRI activity at each different region and their associated predicted values of the g-factor, using the *effects* package (Fox and Weisberg 2018). For multivariate algorithms, we plotted Accumulated Local Effects (ALE) (Apley and Zhu 2020). ALE is defined as follows:

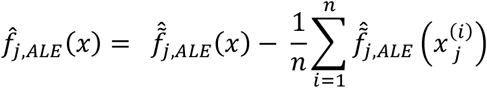

where

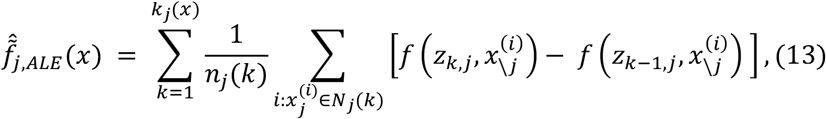

First, ALE creates a grid, z, that splits the values of an explanatory variable, j, into small windows. The difference between predicted values of x_j_ at the two edges of each window is the ‘effect’. The summation of the effects across data points within the same window, k, is the ‘local effect’. The accumulation of local effects from the first window to the last window then constitutes ‘accumulated local effects, ALE’. Finally, the value of ALE is mean centred, such that the main effect of the explanatory variable at a certain value is compared to the average prediction (Molnar 2019). Accordingly, with ALE, we could plot a line to show how a brain region impacted the prediction of each algorithm on average. We computed ALE with a grid size of 20 using the *FeatureEffect* command from the *iml* package (https://christophm.github.io/iml/).

With univariate effects and ALE, we examined the similarity in the pattern and directionality across algorithms. We picked the top 30 regions with the highest variable importance across algorithms. More specifically, we were interested to see which multivariate algorithms provided the pattern and directionality similar to those of univariate algorithms.

#### Interaction: Friedman’s H Statistic

Some algorithms, including Random Forest, XGBoost and SVM with RBF and polynomial kernels, allow for interactions among explanatory variables. Here, we used Friedman’s H-statistic (Friedman and Popescu 2008) to reveal interaction strength from different brain regions. Friedman’s H-statistic relies on a partial dependent (PD) function (Friedman 2001):

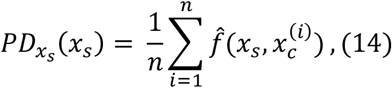

where *x_s_* is the explanatory variable of interest, and 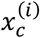 are actual values for all other explanatory variables. Accordingly, PD indicates the average marginal effect for a given value of the explanatory variable *x_s_* (Molnar 2019). Friedman’s H-statistic, in turn, is estimated from

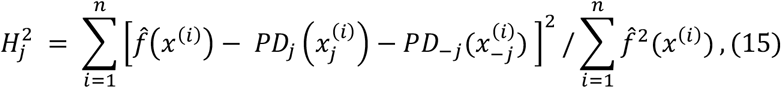

where H is Friedman’s H-statistic. Here 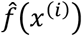 is the prediction function with all explanatory variables included. 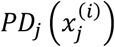 and 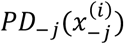 are PD for an explanatory variable j, and for those without the explanatory variable j, respectively. In the case of no interaction, 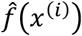 is the same with the sum of 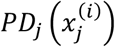 and 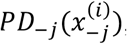, which results in Friedman’s H-statistic of 0. Friedman’s H-statistic of 1 means that the variation of the prediction is only due to interaction. Accordingly, a higher Friedman’s H-statistic indicates a higher interaction strength from a certain explanatory variable. We computed Friedman’s H-statistic using the *Inleradion$new()* command from the *iml* package (https://christophm.github.io/iml/) and plotted 20 brain regions with the highest Friedman’s H-statistic from each algorithm.

Please see our Github page for our scripts and detailed outputs: https://narunpat.github.io/TaskFMRIEnetABCD/ExplainableMachineLearningForTaskBasedfMRI.html

## Results

### The g-factor

Figure 1 shows the confirmatory Factor Analysis (CFA) of the g-factor. The bifactor model of the g-factor shows a good fit: scaled, robust CFI=.992, TLI=.983 and RMSE= .028 (90%CI=.021- .035).

**Figure 1.**
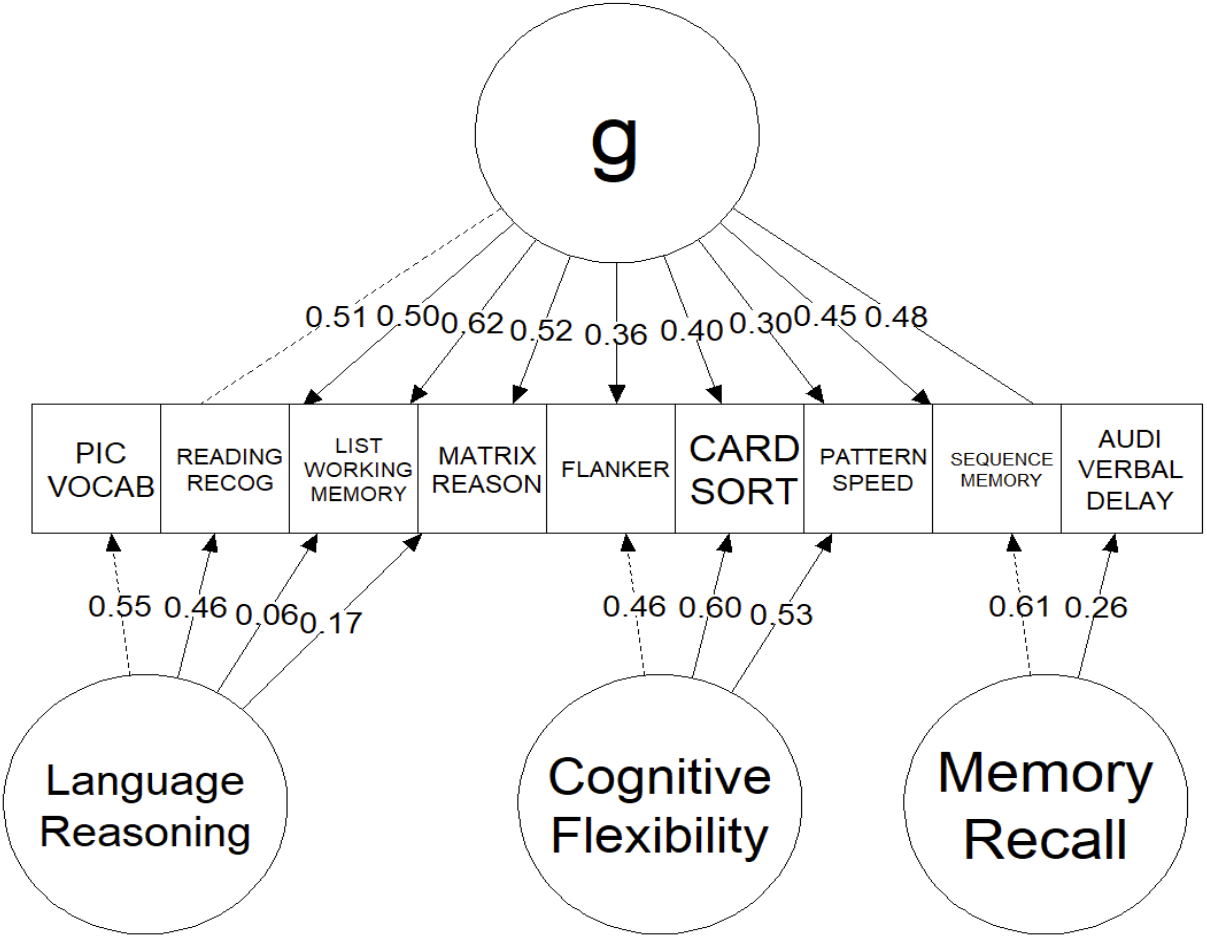
Confirmatory Factor Analysis (CFA) of the bifactor model of the g-factor. The number in each line reflects the magnitude of standardized parameter estimates. The dotted lines indicate marker variables that were fixed to 1. Pic Vocab = Picture Vocabulary; Reading Recog = Oral Reading Recognition; Pattern Speed = Pattern Comparison Processing task.

### Predictive performance

Figure 2 shows the bootstrap distributions of predictive performance across algorithms and response variables. Overall, the mass univariate analyses, either with the FDR or Bonferroni correction, consistently performed worse than multivariate algorithms across response variables and prediction matrices. To statistically compare the predictive ability of linear and additive multivariate algorithms against non-linear and interactive multivariate algorithms along with mass-univariate analyses, we created the bootstrap distributions of the differences in predictive performance by subtracting the performance of the OLS multiple regression (Figure 3, Supplementary Figure 1) and Elastic Net (Figure 4, Supplementary Figure 2) from other algorithms. Note to highlight the differences between linear and additive vs. non-linear and interactive multivariate algorithms, we created Supplementary Figures 1 and 2 the comparisons without mass univariate analyses. These comparisons revealed problems with the OLS multiple regression. For most response variables, the predictive performance of the OLS multiple regression was significantly worse than most of the machine-learning algorithms. Moreover, when the response variables were not-well predicted (e.g., sequential memory, Flanker, auditory-verbal and pattern speed), the predictive performance of the OLS multiple regression was even worse than that of mass univariate analyses. By contrast, across the response variables, the performance of Elastic Net was either on par with or better than other algorithms. Lastly, fMRI during the n-back task predicted the g-factor well, compared to other out-of-scanner cognitive tasks, especially with multivariate algorithms.

**Figure 2.**
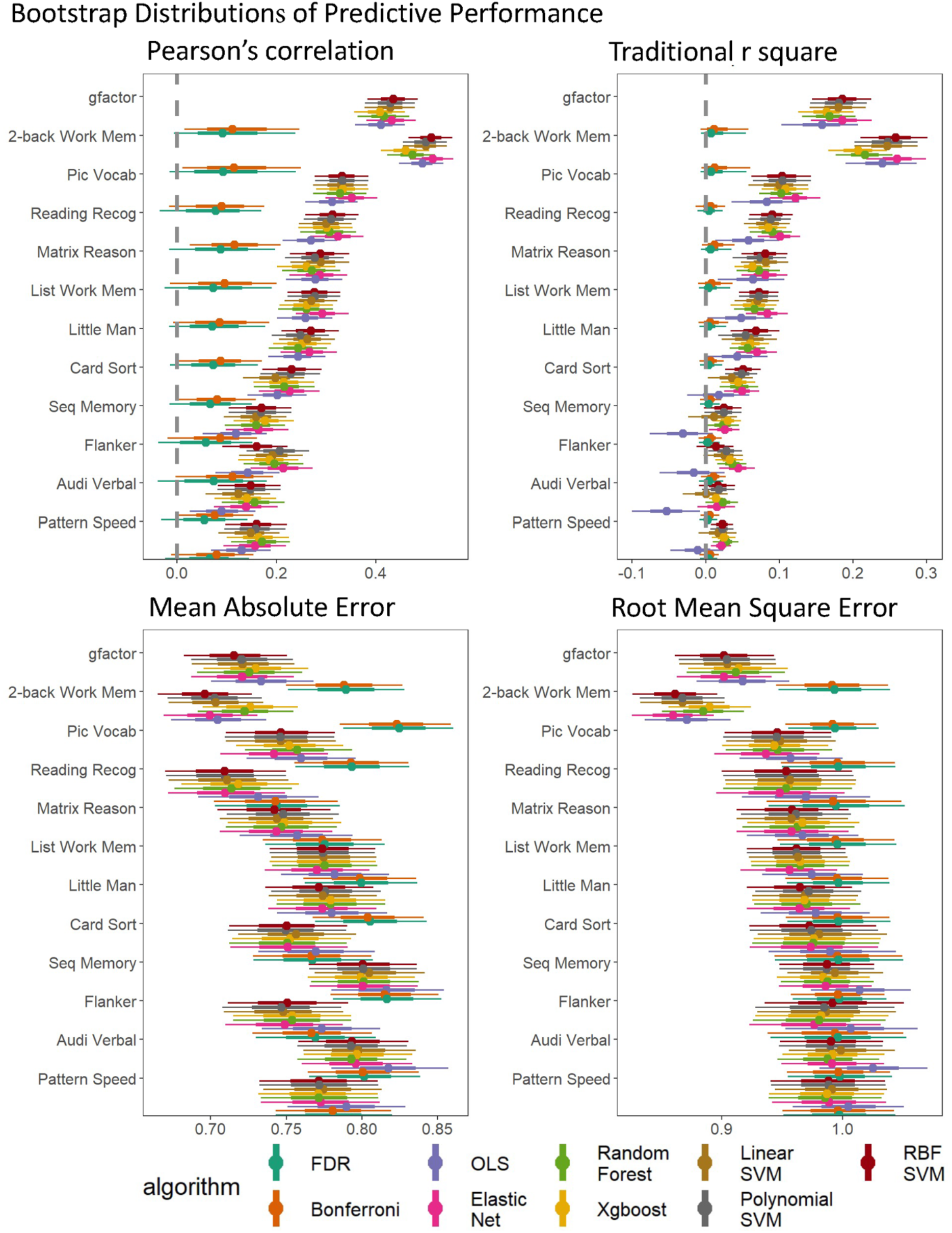
Bootstrap distributions of predictive performance across algorithms and response variables. The thicker lines reflect 65% bootstrap confidence intervals and the thinner lines reflect 95% confidence intervals. MAE = the mean absolute error; RMSE = root mean square error.

**Figure 3.**
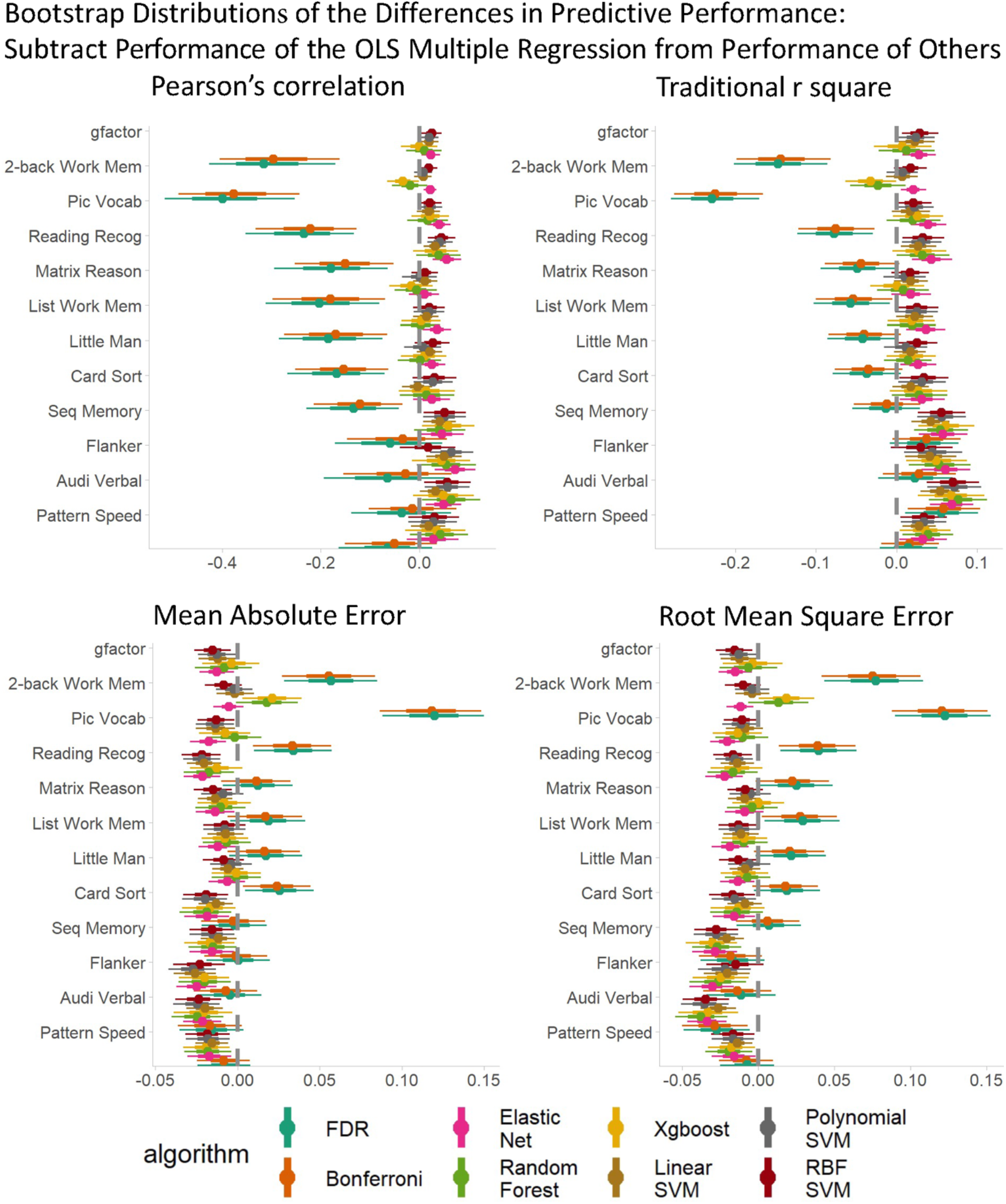
Bootstrap distributions of the differences in predictive performance between the OLS multiple regression and other algorithms. For Pearson’s correlation and traditional r square, values lower than zero indicates lower performance than the OLS multiple regression. For mean absolute error and root mean square error, values higher than zero indicates lower performance than the OLS multiple regression. The thicker lines reflect 65% bootstrap confidence intervals and the thinner lines reflect 95% confidence intervals. MAE = mean absolute error; RMSE = root mean square error.

**Figure 4.**
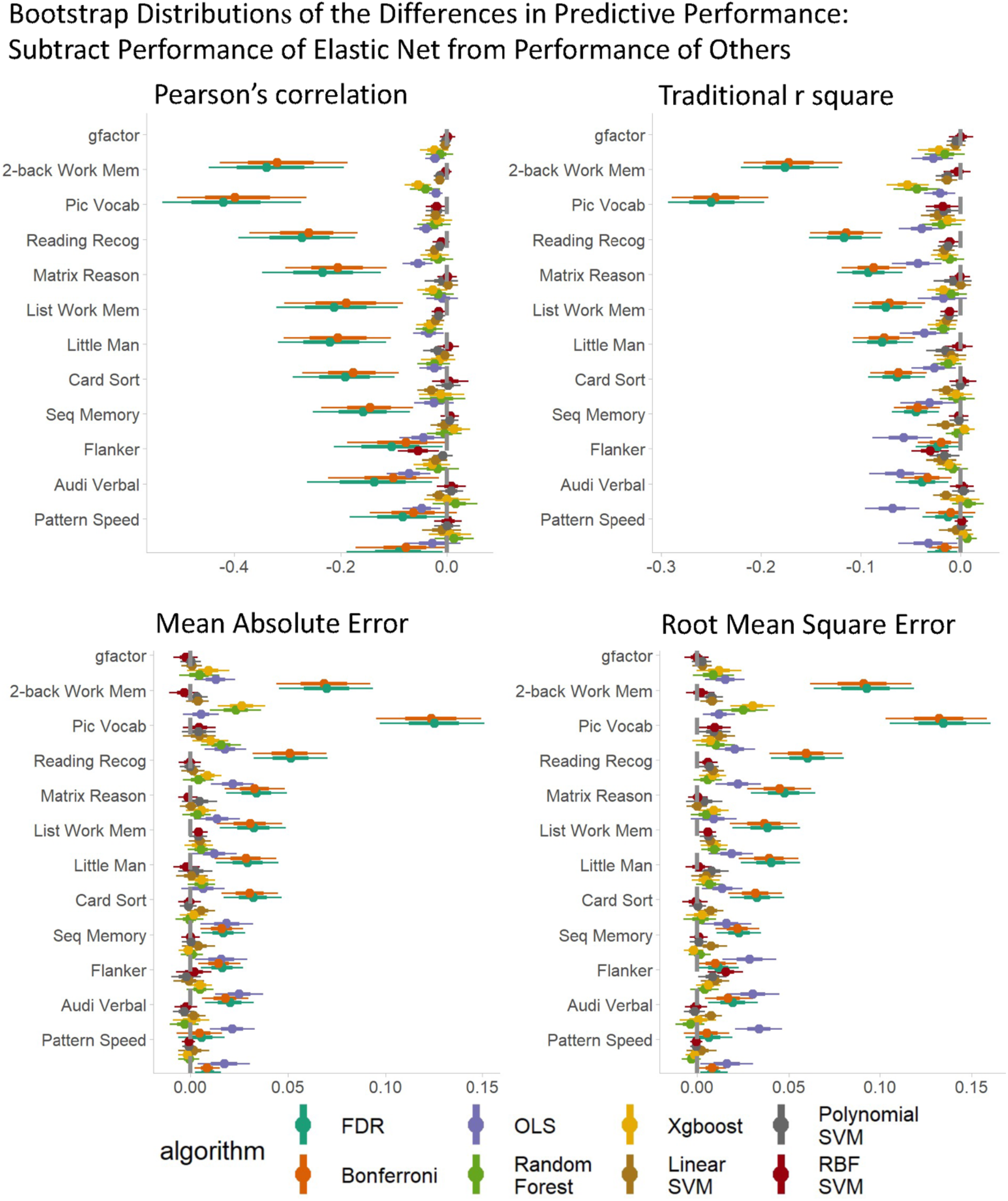
Bootstrap distributions of the differences in predictive performance between Elastic Net and other algorithms. For Pearson’s correlation and traditional r square, values lower than zero indicates lower performance than Elastic Net. For mean absolute error and root mean square error, values higher than zero indicates lower performance than Elastic Net. The thicker lines reflect 65% bootstrap confidence intervals and the thinner lines reflect 95% confidence intervals. MAE = mean absolute error; RMSE = root mean square error.

### Explaining the algorithms

#### Variable importance: Coefficients and SHAP

Figure 5 shows the variable importance of models predicting the g-factor across all algorithms. Figure 6 shows Spearman’s correlations in variable importance among the algorithms. The variable importance of different algorithms appear to be significantly related (i.e., ρ with p-value < .05), but in varying degrees. For instance, the variable importance of Elastic Net was more similar to that of SVM of different kernels (ρ ~ .7) than that of mass-univariate algorithm (ρ ~.2). On the other hand, the variable importance of mass-univariate algorithms seemed more closely related to that of Random Forest and XGBoost (ρ ~.5-.6) than other algorithms. Accordingly, this shows that different algorithms drew information from areas across the brain differently.

**Figure 5.**
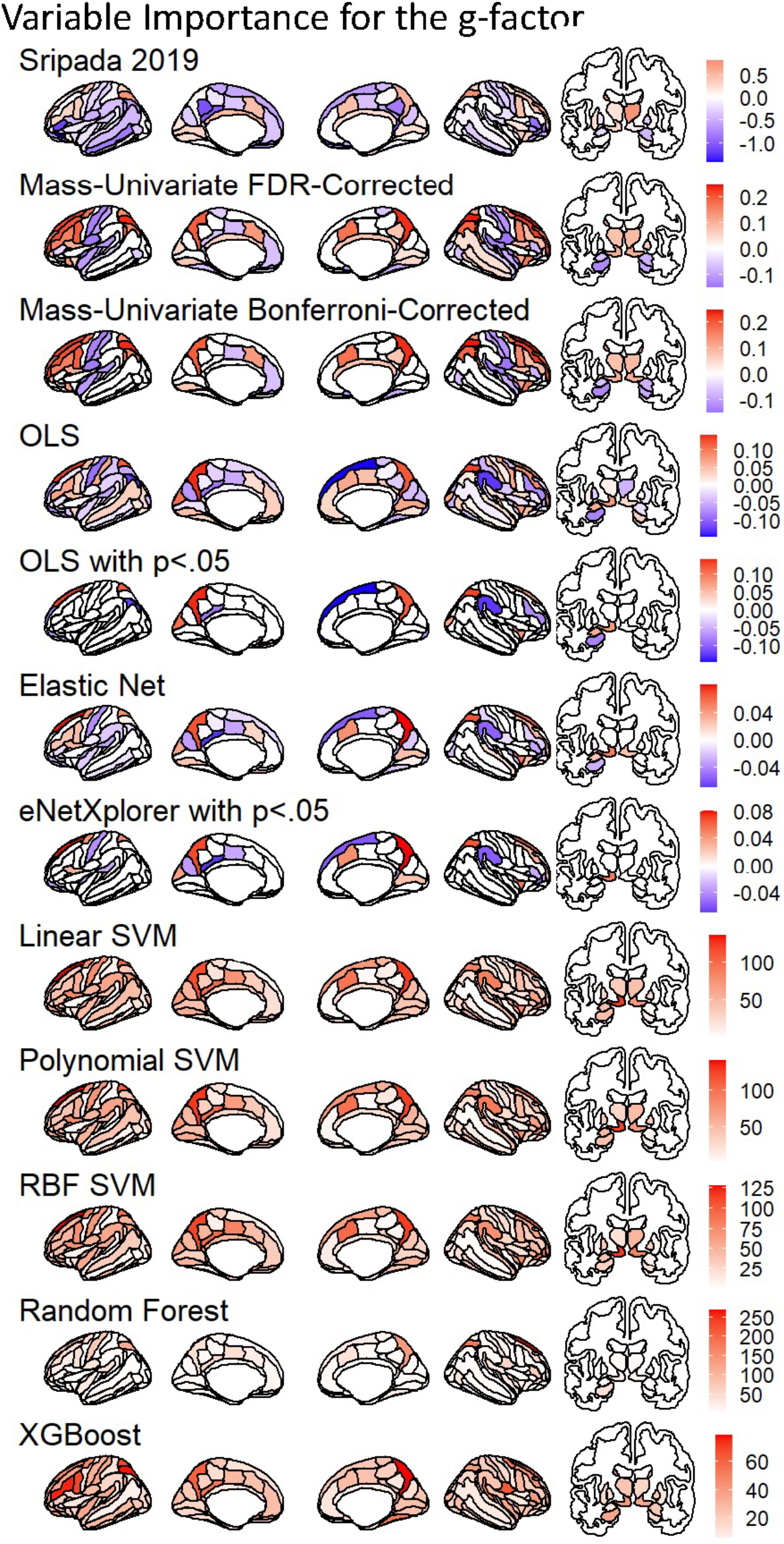
Variable importance of models predicting the g-factor across all algorithms. Variable importance for Sripada et al., (2019) is based on a weighted, principal component regression. Variable importance for Mass Univariate Analyses, OLS multiple regression and Elastic Net is the coefficient values while variable importance for SVM, Random Forest, XGBoost is the absolute value of SHAP. We plotted variable importance on the brain using ggseg (Mowinckel and Vidal-Piñeiro 2019).

**Figure 6.**
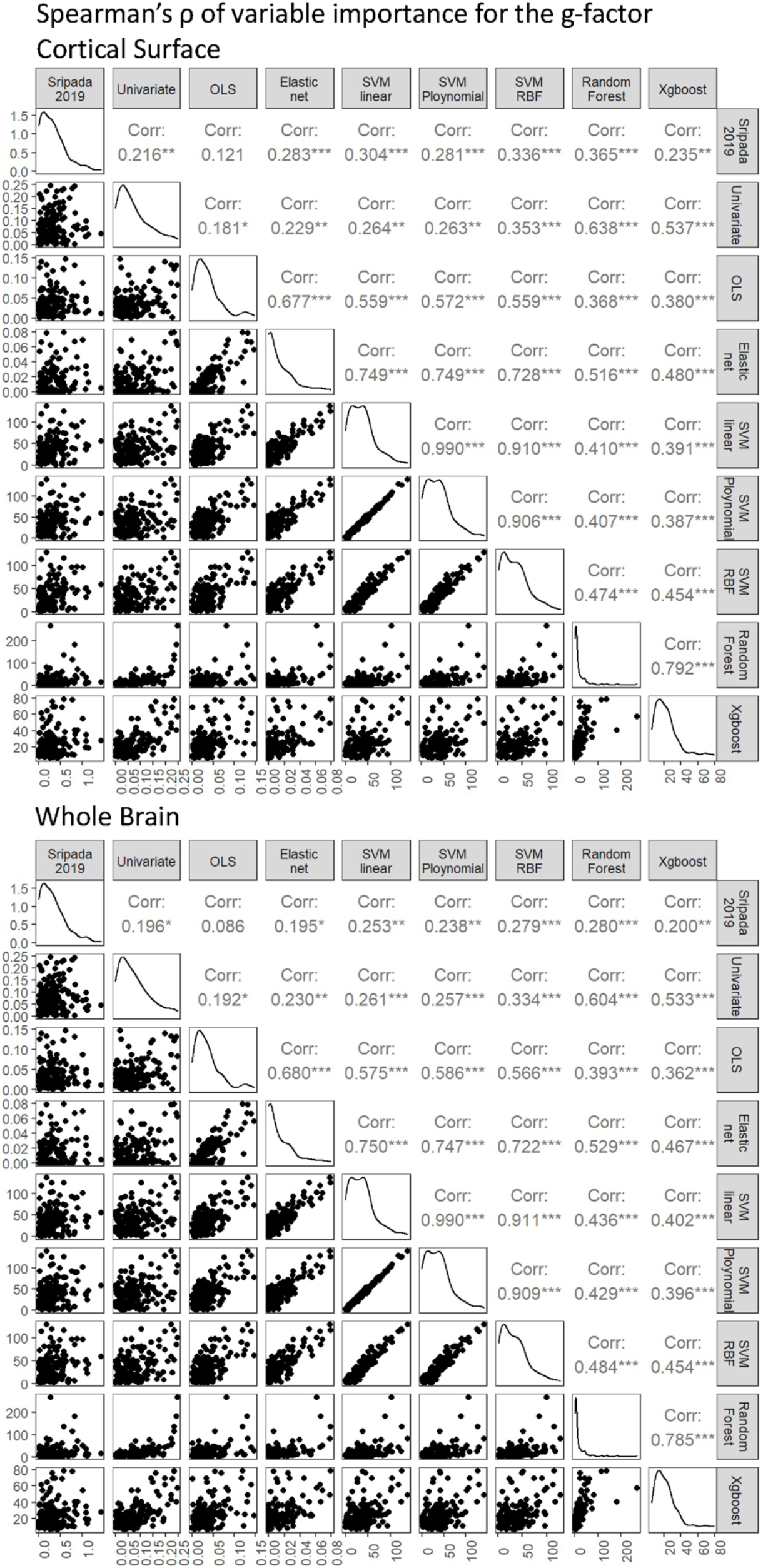
Spearman’s correlations in variable importance between algorithms. We calculated variable importance for models predicting the g-factor. Variable importance for Sripada et al., (2019) is based on a weighted, principal component regression. Variable importance for Mass Univariate Analyses, OLS multiple regression and Elastic Net is the coefficient magnitude (i.e., the absolute value of the coefficients). Variable importance for SVM, Random Forest, XGBoost is the absolute value of SHAP.

As for the similarity with Sripada and colleagues’ (2020) findings, we found significant Spearman’s correlations (p-value <.05) across all but one algorithm, the OLS multiple regression. These significant correlations were small in magnitude (ρ ~.2-.3 on the cortical surface and ρ ~.2 on the whole brain). Still, the OLS multiple regression seemed to be the only algorithm that yielded inconsistent results with Sripada and colleagues’ (2020) findings.

#### Variable selection for variable importance: Conventional p-value and eNetXplorer

Figure 5 plots variable selection for the variable importance of models predicting the g-factor on the brain. These plots include mass univariate analyses with FDR and Bonferroni corrections, the OLS multiple regression, and Elastic Net with eNetXplorer. Figures 7 plots variable selection for the OLS multiple regression and Elastic Net to highlight the differences between the OLS multiple regression and eNetXplorer in coefficient magnitude and uncertainty estimates as a function of hyperparameters of Elastic Net and multicollinearity. The OLS multiple regression selected 23 regions while eNetXplorer selected 27, 32 and 21 regions for full Elastic Net, Ridge and LASSO, respectively (Figure 7, 8). Of these selected regions, only 14 regions were the same regions among the OLS multiple regression, full Elastic Net, Ridge and LASSO, and 20 regions were the same among Elastic Net, Ridge and LASSO. Compared to Elastic Net (penalty at .13) and LASSO (penalty at .01), Ridge led to the highest penalty at .36, resulting in the smallest coefficient magnitude. As for uncertainty estimated, we found that, for the OLS multiple regression, the coefficient SE linearly increased as a function of the VIF. In contrast, for eNetXplorer, the changes in the permuted null coefficient SD as a function of the VIF were less pronounced.

**Figure 7.**
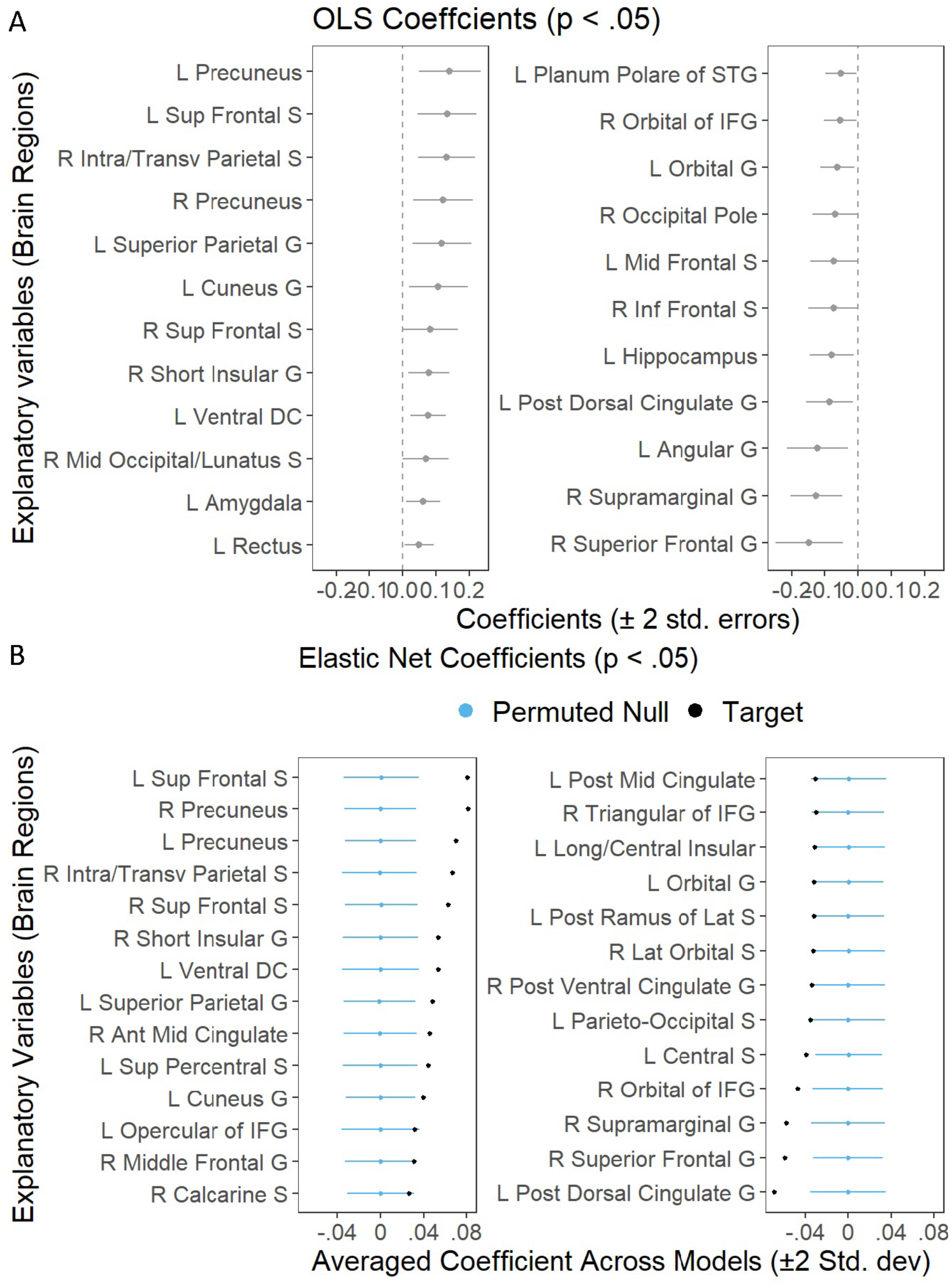
Variable selection for the variable importance of the OLS multiple regression (A) and Elastic Net (via eNetXplorer) (B) predicting the g-factor. We only plotted regions with p<.05.

**Figure 8.**
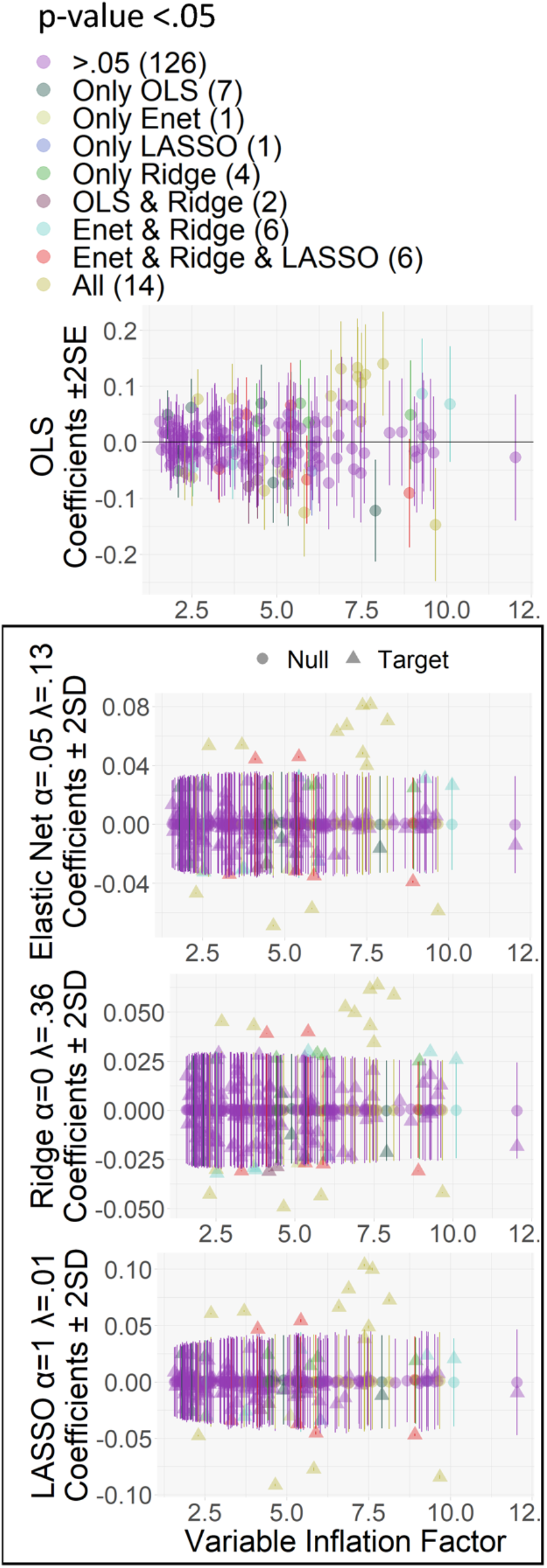
Uncertainty estimates of explanatory variables (brain regions) as a function of Variance Inflation Factor (VIF) for the OLS multiple regression, Elastic Net, Ridge and LASSO predicting the g-factor. We calculated the VIF based on the OLS multiple regression. For the OLS multiple regression, confidence intervals are the coefficients plus/minus 2 multiplied by coefficient’s standard error (SE). For the Elastic Net, Ridge and LASSO, confidence intervals are the permuted null coefficients plus/minus 2 multiplied by the permuted null standard deviation (SD).

#### Prediction pattern and directionality: Univariate Effects and Accumulated Local Effects

Figure 9 plots univariate effects and Accumulated Local Effects (ALE) of models predicting the g-factor across all algorithms. They show the prediction pattern and directionality of the relationship between predictive values of the g-factor based on different algorithms and fMRI activity at each brain region. Across brain regions, most of the multivariate algorithms, apart from the OLS multiple regression, had a similar pattern and direction a) with each other and b) with the univariate effects. ALE of the OLS multiple regression deviated from that of other multivariate algorithms in many regions. Moreover, ALE of the OLS multiple regression demonstrated a relationship in an opposite direction to univariate effects in several regions, such as the left intra-transverse parietal sulcus, left middle frontal gyrus, left and right middle frontal sulcus, right supplementary precentral sulcus, left angular gyrus, right post ramus of the lateral sulcus and right superior parietal gyrus.

**Figure 9.**
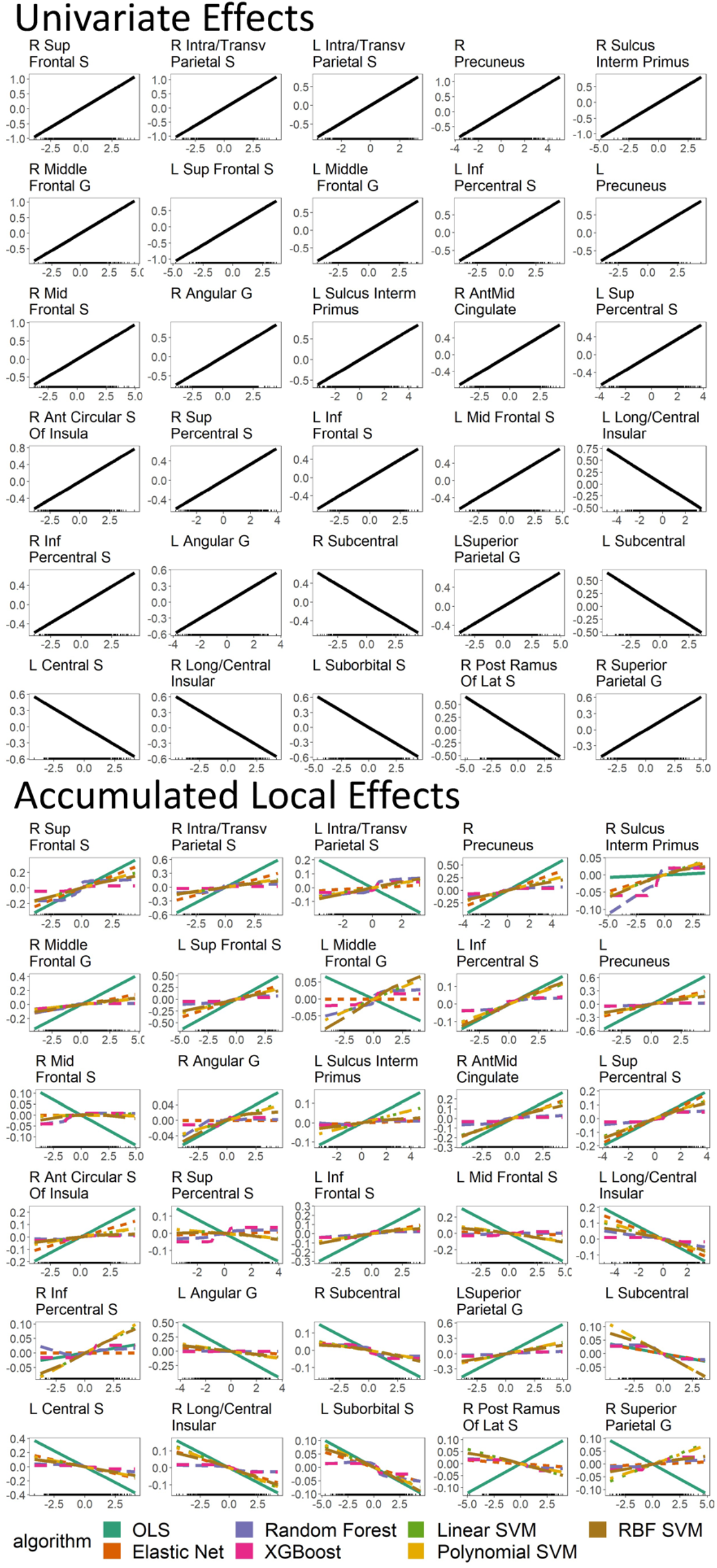
Univariate effects and Accumulated Local Effects (ALE) of models predicting the g-factor across all algorithms. These plots show the prediction pattern and directionality of the relationship between predictive values of the g-factor from different algorithms and fMRI activity at each brain region. We plotted the top 30 brain regions with the highest variable importance across algorithms.

#### Interaction: Friedman’s H Statistic

Figure 10 show interaction plots based on Friedman’s H-statistic. They show the interaction strength between each brain region and all other brain regions for four interactive algorithms predicting the g-factor, including Random Forest, XGBoost and SVM with RBF and polynomial kernels. The interaction strength between brain regions for XGBoost and SVM with polynomial and RBF kernels accounted for less than 6% of variance explained per region. Random forest, on the other hand, had two explanatory variables that accounted for 15% and 20% of variance explained per region.

**Figure 10.**
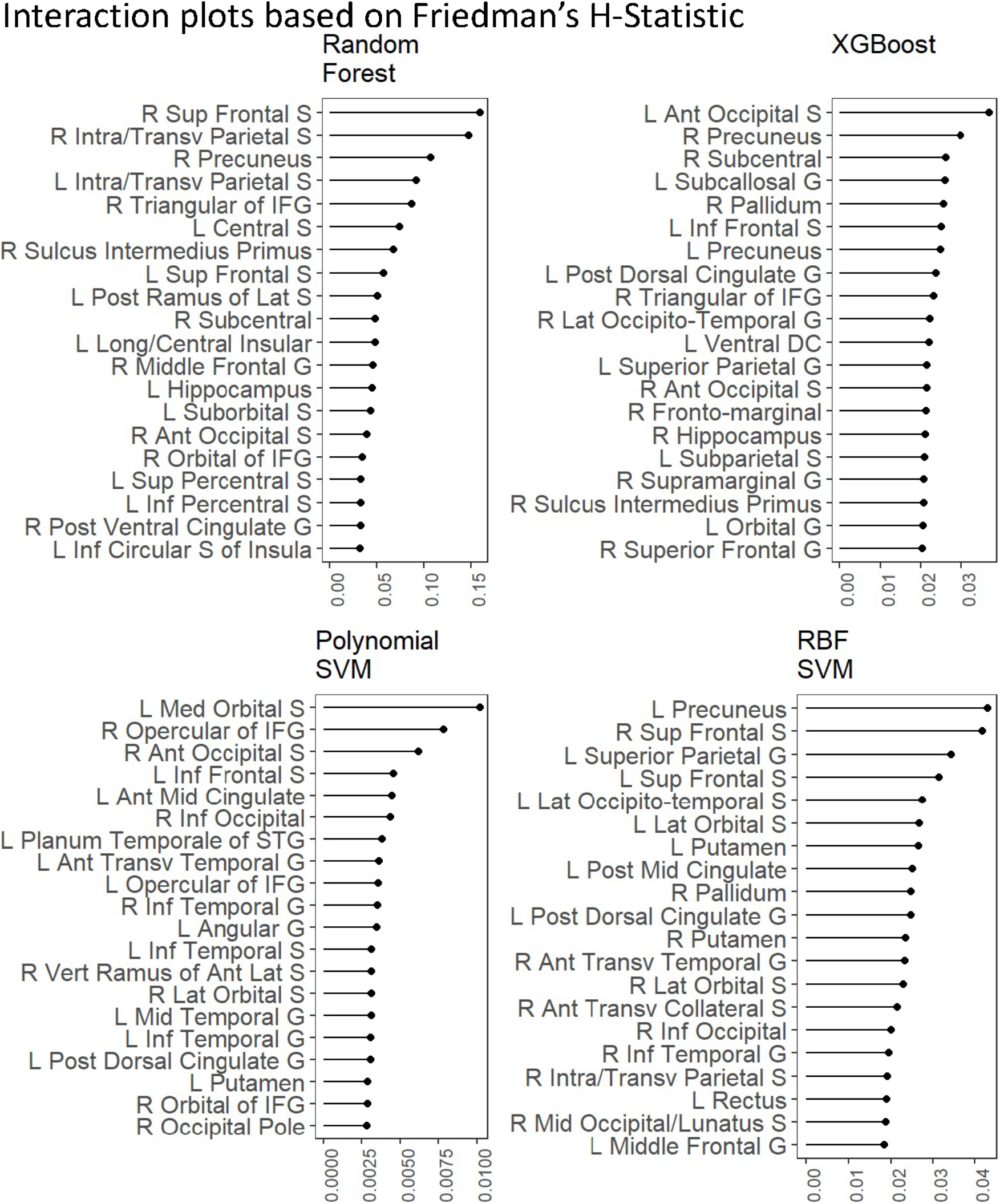
Interaction plots based on Friedman’s H-statistic. H-statistic here indicates the interaction strength between each brain region and all other brain regions for different algorithms predicting the g-factor. Higher values indicate higher strength of the interaction from a particular brain region. Here we plotted the top 20 brain regions with the highest Friedman’s H-statistic at each algorithm.

## Discussion

We applied the explainable machine-learning framework (Molnar 2019; Belle and Papantonis 2021) to predict children’s cognition from task-based fMRI during the n-back task, using the ABCD dataset (Casey et al. 2018). We first compared the performance of nine algorithms in their ability to predict individual differences in cognitive abilities across 12 response variables and found better predictive performance from machine learning algorithms, compared to the conventional mass-univariate analyses and the OLS multiple regression. Despite being a linear and additive algorithm, Elastic Net came up among the top-performing algorithm. We then implemented various explainers to explain how these algorithms drew information from task-based fMRI to predict the g-factor. With these explainers, we found 1) some similarity in variable importance across algorithms and studies, 2) differences in variable selection between the OLS multiple regression and Elastic Net, 3) similar directionality in the relationship with the g-factor between machine-learning and mass-univariate algorithms, and 4) interaction from certain brain regions as captured by interactive algorithms. These explainers also showed potential problems of the OLS multiple regression, including having lower consistency in variable importance with a prior study (Sripada et al. 2020) and having a relationship with the g-factor in the opposite direction with the mass-univariate algorithms at many regions, suggesting the presence of suppression (Ray-Mukherjee et al. 2014).

For predictive performance, the conventional mass-univariate algorithms, either with the FDR or the more-conservative Bonferroni correction, showed worse performance than most multivariate algorithms across most response variables of individual differences in cognitive abilities. Our findings using task-based fMRI are consistent with recent work that compared the performance between mass-univariate and multivariate algorithms (via SVM with the RBF kernel) to predict individual differences in cognition from resting-state fMRI and structural MRI (Marek et al. 2022). Accordingly, across different MRI modalities, using multivariate algorithms appear to be a more promising approach than the mass-univariate algorithms to ensure reproducibility of the brain-cognition relationship and to use MRI as a predictive tool for individual differences.

Nonetheless, simply including all brain regions as explanatory variables in an OLS multiple regression model may not be ideal for prediction. The OLS multiple regression had poorer predictive performance than machine-learning algorithms for most response variables. More importantly, its predictive performance for response variables that were hard to predict across algorithms (e.g., sequential memory, Flanker, auditory-verbal and pattern speed) was very poor, indicated by negative traditional r square and lowest RMSE, even when compared to the mass-univariate algorithms. By contrast, adding regularisation to the OLS multiple regression algorithm in the form of Elastic Net (Zou and Hastie 2005) appeared to boost predictive performance across response variables. In fact, the performance of Elastic Net, despite being a less complex machine-learning algorithm given its constraints on linearity and additivity, was on par with, and in many cases better than, many nonlinear and interactive algorithms, including Random Forest, XGBoost and SVM of different kernels. This finding is consistent with work on resting-state fMRI, showing that a penalized regression-based algorithm, such as Elastic Net, performed just as well as other algorithms (Dadi et al. 2019). Given that Elastic Net is readily interpretable, it can be considered a parsimonious choice for future task-based fMRI studies.

We observed variability in the level of predictive performance among the 12 response variables of individual differences in cognitive abilities, especially with multivariate algorithms. It is not surprising to see the highest predictive performance from the behavioural performance during the n-back fMRI task. Given that the fMRI and behavioural data were collected at the same time, our predictive models may capture idiosyncratic variation due to the task and session itself (e.g., arousal, attention and other processes). As for out-of-scanner tasks, our predictive models performed better for some cognitive tasks (e.g., picture vocabulary, reading recognition and matrix reasoning) than others (e.g., sequential memory, Flanker, auditory-verbal and pattern speed). More importantly, capturing the shared variance of behavioural performance across different tasks using the latent variable, the g-factor, led to higher predictive performance than any individual task. This suggests that our framework can be used to build predictive models for individual differences in cognitive abilities in general, and is not confined to specific tasks or processes.

Our implementation of various explainers allowed us to better understand how each algorithm made a prediction of the g-factor. First, variable importance enabled us to move beyond simply estimating the contribution from each separate brain region (i.e., marginal importance), commonly done in mass-univariate analyses. Instead, with variable importance, we could investigate the contribution from multiple regions in the same model (i.e., partial importance) (Chen et al. 2019; Debeer and Strobl 2020). Here, we found different degrees of similarity in variable importance across algorithms. On the one hand, the variable importance of mass-univariate algorithms, which are the dominant approach for explaining the brain-cognition associations in the neuroimaging community (Friston 2007), were more closely related to tree-based algorithms, Random Forest and XGBoost, than other algorithms. On the other hand, the top two algorithms that predicted the g-factor well in the current study, Elastic Net and SVM with the RBF kernel, had variable importance that was correlated well with each other (ρ > .7). This suggests that the brain information similarly drawn by Elastic Net and SVM with the RBF kernel in our study allowed us to capture individual differences in the g-factor relatively well.

More importantly, we also tested the similarity in variable importance found in the current study with Sripada and colleagues’ (2020). Variable importance from all algorithms, except for the OLS multiple regression, was significantly correlated with that of Sripada and colleagues (2020). While significant, these correlations were small in magnitude, perhaps due to many different characteristics between Sripada and colleagues’s (2020) and our study. For instance, Sripada and colleagues (2020) built a predictive model from adults’ data (as opposed to children’s data), applied principle component regression on non-parcellated regions (as opposed to different univariate and multivariate algorithms on parcellated regions) and used a slightly different variation of the n-back task (Barch et al. 2013; Casey et al. 2018) and of the response variables for individual differences in cognitive abilities. Additionally, we also saw more similarity in variable importance with Sripada and colleagues (2020) on the cortical surface, compared to the whole brain. This might be due to the superiority of cortical surface in brain registration across ages, compared to the subcortical volumetric regions used in the whole brain (Ghosh et al. 2010). Altogether, this suggests some degree of consistency in variable importance across studies for most of the algorithms examined here, apart from the OLS regression.

We showed variable selection for different algorithms: mass univariate analyses with FDR and Bonferroni corrections, the OLS multiple regression, and Elastic Net with eNetXplorer (Candia and Tsang 2019). Being able to explain and select brain regions that exhibited an above-chance-level contribution to the prediction of Elastic Net via eNetXplorer was of great importance, given that Elastic Net had high predictive performance in the current study. Variable selection for the OLS multiple regression and eNetXplorer relies on two factors: coefficient magnitudes and uncertainty estimates. Accordingly, we created Figure 8 to investigate coefficient magnitudes and uncertainty estimates at different levels of the mixture hyperparameter, including tuned (Elastic Net) or fixed at either 0 (Ridge) or 1 (LASSO), and multicollinearity. For coefficient magnitudes, we saw lower coefficient magnitude from an algorithm with higher regularisation: ranking from Ridge, (penalty (λ) = .36), Elastic Net (penalty = .13), LASSO (penalty = .01) to the OLS multiple regression, which could be viewed as having penalty at 0. As for uncertainty estimates, we saw two behaviours of eNetXplorer. First, in our data, the permutation used in eNetXplorer showed more consistency in uncertainty estimates across the different levels of multicollinearity (as reflected by VIF), as compared to the OLS multiple regression. Second, a higher regularisation, indicated by the penalty, led to not only a smaller coefficient magnitude of the target model, but also a smaller standard deviation (SD) of the permuted null model, which is the uncertainty estimate for eNetXplorer. This is because eNetXplorer used the same hyperparameters for both the target and permuted null models, which were based on the hyperparameters associated with the best predictive performance for the target model during cross-validation. eNetXplorer then tested which of the brain regions contributed to the prediction of regularised regression higher than chance by comparing regularised coefficients of the target models against regularised coefficients of the permuted null models. Thus, a higher reduction in coefficient magnitude based on the penalty hyperparameter does not necessarily mean a smaller number of regions selected. To illustrate this, in our case, Ridge led to the highest penalty, which resulted in the smallest coefficient magnitude, but had the highest number of regions selected. Altogether, as a result of Elastic Net hyperparameters and multicollinearity, some of the regions with high VIF had high SE and were not selected by the OLS multiple regression, but were selected by eNetXplorer. In contrast, some of the regions that were selected by the OLS multiple regression had smaller relative magnitude and were not selected by eNetXplorer. Thus, eNetXplorer allowed us to explain and select variables that contributed to the prediction given the combination of Elastic Net hyperparameters estimated (Candia and Tsang 2019).

Next, to reveal the pattern and directionality of the relationship between task-based fMRI activity and the g-factor based on different algorithms at different brain regions, we used univariate effects and Accumulated Local Effects (ALE). For the pattern, ALE revealed the expected pattern from each algorithm. For instance, ALE showed a linear pattern from linear algorithms, such as the OLS multiple regression and Elastic Net. Similarly, ALE also showed a non-linear pattern from non-linear algorithms, such as Random Forest and XGBoost. As for the directionality, we saw inconsistency in the directionality between ALE from the OLS multiple regression and univariate effects from mass-univariate analyses. For example, there were many brain regions that were shown to have a positive relationship according to univariate effects, but have a negative relationship according to ALE from the OLS multiple regression. Some of these regions included the left intra/transverse parietal sulcus, left middle frontal gyrus, left and right middle frontal sulcus, right supplementary precentral sulcus, left angular gyrus and right superior parietal gyrus. Accordingly, adding multiple brain regions into an OLS multiple-regression model appeared to change the directionality of the relationship each of the regions had when considered by itself. This may indicate the “suppression” effect whereby the directionality of an explanatory variable depends on its relationship with other explanatory variables (Courville and Thompson 2001; Beckstead 2012; Ray-Mukherjee et al. 2014). Fortunately, all of the machine-learning algorithms we tested did not show this inconsistency. Indeed, regularising the coefficients in the OLS multiple regression as implemented by Elastic Net appeared to keep the directionality of the relationship in line with that of mass-univariate analyses.

Finally, using Friedman’s H-statistic (Friedman and Popescu 2008), we also demonstrated the interactions among brain regions that were captured by each of the four interactive algorithms: Random Forest, XGBoost and SVM with polynomial and RBF kernels. The interaction strength between brain regions for XGBoost and SVM with polynomial and RBF kernels was quite weak, accounting for less than 6% of variance explained per region. Random forest, on the other hand, had two brain regions, the right superior frontal sulcus and right intraparietal sulcus, that accounted for 15% and 20% of variance explained per region. Nonetheless, these four algorithms that allowed for interactions generally did not perform well over non-interactive Elastic Net in terms of predictive performance for the g-factor. This means that not accounting for interactions may be parsimonious enough for the current data.

Given the emergence of large-scale, task-based fMRI studies (Van Horn and Toga 2014), one potential use of the explainable machine-learning framework (Molnar 2019; Belle and Papantonis 2021) is to build explainable, predictive models for future studies. This is similar to polygenic scores in genomics (Torkamani et al. 2018) where geneticists use large discovery datasets to build models that reflect the influences of SNPs across the genome on certain phenotypes of interest in the form of polygenic scores. Polygenic scores improve the replicability of genetics studies, compared to the classical “candidate-gene” approach (Bogdan et al. 2018). Polygenic scores are explainable based on the influences of each SNP associated with phenotypes in the discovery dataset (Torkamani et al. 2018). Here we suggest that neuroimagers can take a similar approach to build a predictive model based on task-based fMRI from large-scale studies. In fact, our predictive performance for the g-factor from just one fMRI n-back task (r square ~ .2) was considerably higher than that from a polygenic score (r square < .1) (Allegrini et al. 2019). This encourages the potential use of task-based fMRI for individual differences in cognition.

To build predictive models for individual differences in cognition from task-based fMRI, there are existing toolboxes, particularly designed for neuroimaging, such as PRoNTO (Schrouff et al. 2013), pyMVPA (Hanke et al. 2009), and Neurominer (Hanke et al. 2009). Here, we took a more general machine-learning approach using the *tidymodels* (Kuhn et al. 2020) and other R packages. Given the availability of a number of algorithms in *tidymodels* and the versatility of the R language, our approach is more flexible and readily scalable. Moreover, unlike the neuroimaging toolboxes, using the R language allowed us to integrate the predictive modelling framework with modern explainers, such as SHAP (Lundberg and Lee 2017), eNetXplorer (Candia and Tsang 2019), ALE (Apley and Zhu 2020) and Friedman’s H-statistic (Friedman and Popescu 2008). These explainers are not likely to be available in the purposefully built toolboxes for neuroimaging.

Our study has limitations. While the ABCD dataset is one of the largest datasets with task-based fMRI available (Casey et al. 2018), its sample as of 2020 has, by design, a narrow age range. The predictive model built from our study may only be generalized to children aged 9-10 years old. However, given that the study will trace the participants until they are 19-20 years old, future studies will be able to use a similar approach to ours and expand the age range. Next, our method shown relied on Freesurfer’s parcellation (Fischl et al. 2002; Destrieux et al. 2010), which is the only parcellation available post-processed from the ABCD release 2.0.1 (Yang and Jernigan 2019). This commonly used parcellation is based on subject-specific anatomical landmarks, but its regions are relatively large. Future studies may need to demonstrate predictive ability with smaller parcels, which will lead to more regions, and in turn, more explanatory variables, but might also result in higher inconsistencies in identifying areas across participants. Lastly, there were many large outliers in the data that we dealt with via listwise deletion. Many of our algorithms tested, including the mass univariate, OLS multiple regression, and Elastic Net, all rely on minimizing the sum of the squared errors, which can be disproportionately influenced by outliers (Maronna 2011; Rousselet et al. 2017). In the future, it may be useful to test the workflow that mitigates the influences of outliers without using the listwise deletion (Lawrence and Marsh 1984; Maronna 2011).

To conclude, applying the explainable machine-learning framework (Molnar 2019; Belle and Papantonis 2021) to task-based fMRI from a large-scale study, we demonstrated poorer predictive ability from the conventional mass-univariate approach and the OLS multiple regression. Drawing information from multiple brain areas across the whole brain via different machine-learning algorithms appeared to improve predictive ability. Elastic Net, a linear and additive algorithm, provides predictive performance on par with, if not better than, other algorithms. Using different explainers allowed us to explain different aspects of the predictive models: variable importance, variable selection, pattern and directionality and interaction. This in turn enabled us to pinpoint problems with the OLS multiple regression. We believe our approach should enhance the scientific understanding and replicability of task-based fMRI signals, similar to what polygenic scores have done for genetics.

## Acknowledgments

Data used in the preparation of this article were obtained from the Adolescent Brain Cognitive Development (ABCD) Study (https://abcdstudy.org), held in the NIMH Data Archive (NDA). This is a multisite, longitudinal study designed to recruit more than 10,000 children age 9-10 and follow them over 10 years into early adulthood. The ABCD Study is supported by the National Institutes of Health and additional federal partners under award numbers U01DA041022, U01DA041028, U01DA041048, U01DA041089, U01DA041106, U01DA041117, U01DA041120, U01DA041134, U01DA041148, U01DA041156, U01DA041174, U24DA041123, U24DA041147, U01DA041093, and U01DA041025. A full list of supporters is available at https://abcdstudy.org/federal-partners.html. A listing of participating sites and a complete listing of the study investigators can be found at https://abcdstudy.org/scientists/workgroups/. ABCD consortium investigators designed and implemented the study and/or provided data but did not necessarily participate in analysis or writing of this report. This manuscript reflects the views of the authors and may not reflect the opinions or views of the NIH or ABCD consortium investigators. The authors also thank developers who have maintained R packages used here, including but not limited to eNetXplorer (Candia and Tsang 2019), ggseg (Mowinckel and Vidal-Piñeiro 2019), tidyverse (Wickham et al. 2019) and tidymodels (Kuhn et al. 2020).

NP and YW were supported by Health Research Council Funding (21/618) and by the University of Otago. JC was supported by the Intramural Research Program of the National Institute on Aging, National Institutes of Health (Baltimore, MD, USA).

---

**Supplementary Figure 1.**
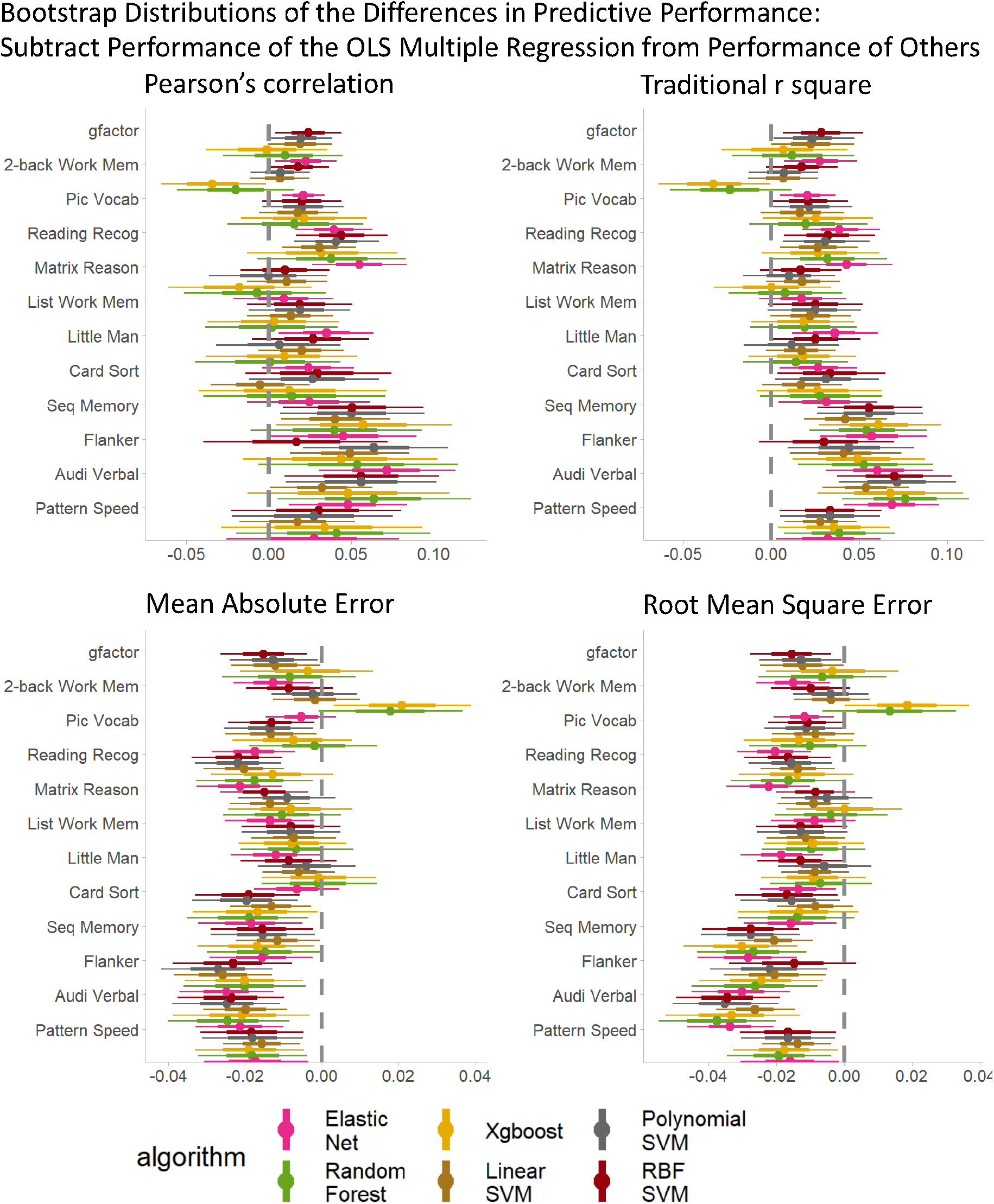
Bootstrap distributions of the differences in predictive performance between the OLS multiple regression and other multivariate algorithms. For Pearson’s correlation and traditional r square, values lower than zero indicates lower performance than <u>the OLS multiple regression</u>. For mean absolute error and root mean square error, values higher than zero indicates lower performance than <u>the OLS multiple regression</u>. The thicker lines reflect 65% bootstrap confidence intervals and the thinner lines reflect 95% confidence intervals. MAE = mean absolute error; RMSE = root mean square error.

**Supplementary Figure 2.**
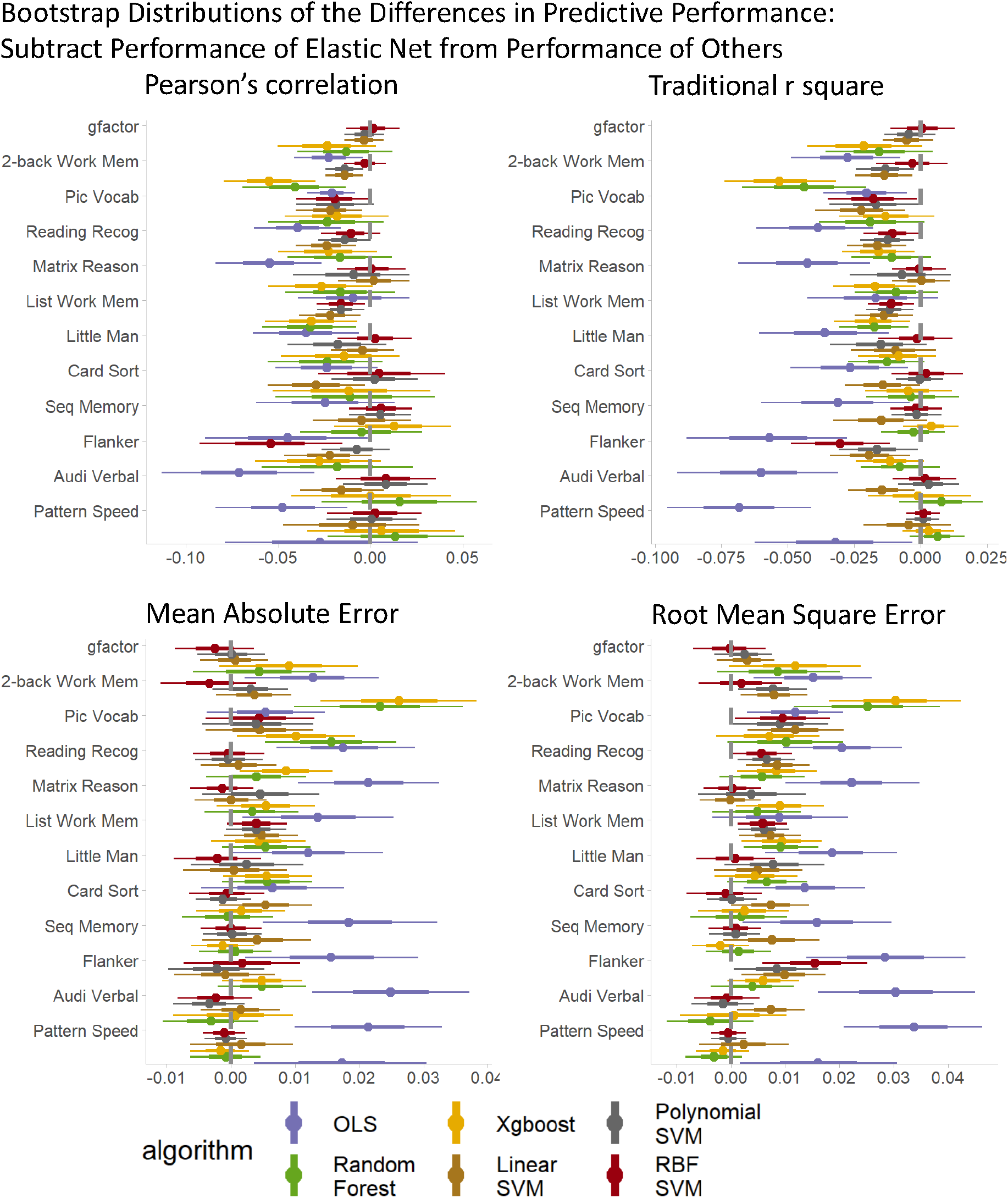
Bootstrap distributions of the differences in predictive performance between Elastic Net and other multivariate algorithms. For Pearson’s correlation and traditional r square, values lower than zero indicates lower performance than Elastic Net. For mean absolute error and root mean square error, values higher than zero indicates lower performance than Elastic Net. The thicker lines reflect 65% bootstrap confidence intervals and the thinner lines reflect 95% confidence intervals. MAE = mean absolute error; RMSE = root mean square error.

